# The Pancreatic Tumor Microenvironment Compensates for Loss of GOT2

**DOI:** 10.1101/2020.08.07.238766

**Authors:** Samuel A. Kerk, Lin Lin, Amy L. Myers, Yaqing Zhang, Jennifer A. Jiménez, Peter Sajjakulnukit, Barbara S. Nelson, Brandon Chen, Anthony Robinson, Galloway Thurston, Samantha B. Kemp, Nina G. Steele, Megan T. Hoffman, Hui-Ju Wen, Daniel Long, Sarah E. Ackenhusen, Johanna Ramos, Xiaohua Gao, Li Zhang, Anthony Andren, Zeribe C. Nwosu, Stefanie Galbán, Christopher J. Halbrook, David B. Lombard, David R. Piwnica-Worms, Haoqiang Ying, Howard C. Crawford, Marina Pasca di Magliano, Yatrik M. Shah, Costas A. Lyssiotis

## Abstract

The tumor microenvironment (TME) in pancreatic ductal adenocarcinoma (PDA) restricts vascularization and, consequently, access to blood-derived nutrients and oxygen, which impacts tumor growth. Intracellular redox imbalance is another restraint on cellular proliferation, yet it is unknown if the TME contributes to the maintenance of redox homeostasis in PDA cells. Here, we demonstrate that the loss of mitochondrial glutamate-oxaloacetate transaminase 2 (GOT2), a component in the malate-aspartate shuttle, disturbs redox homeostasis and halts proliferation of PDA cells in vitro. In contrast, GOT2 inhibition has no effect on in vivo tumor growth or tumorigenesis in an autochthonous model. We propose that this discrepancy is explained by heterocellular pyruvate exchange from the TME, including from cancer associated fibroblasts. More broadly, pyruvate similarly confers resistance to inhibitors of mitochondrial respiration. Genetic or pharmacologic inhibition of pyruvate uptake or metabolism abrogated pyruvate-mediated alleviation of reductive stress from NADH buildup. In sum, this work describes a potential resistance mechanism mediated by metabolic crosstalk within the pancreatic TME. These findings have important implications for metabolic treatment strategies since several mitochondrial inhibitors are currently in clinical trials for PDA and other cancers.

## INTRODUCTION

Cancer cells depend on deregulated metabolic programs to meet their energetic and biosynthetic demands^1–3^. Metabolic therapies aim to preferentially target these dependencies^4^. This approach has shown promise in preclinical models of pancreatic ductal adenocarcinoma (PDA) – one of the deadliest major cancers, notoriously resistant to other modern anti-cancer therapies^5,6^. Pancreatic tumors are poorly vascularized and nutrient austere^7^. Therefore, cancer cells commandeer metabolic pathways to scavenge and utilize nutrients^6,8^. A wealth of recent literature has identified that this is mediated predominantly by mutant KRAS (KRAS*), the oncogenic driver in most pancreatic tumors^9–14^. KRAS* has also been implicated in shaping the pancreatic tumor microenvironment^15,16^. PDA tumors exhibit a complex tumor microenvironment^17,18^ with metabolic interactions between malignant and stromal or immune cells enabling and facilitating tumor progression^19^. Despite recent progress in therapeutic development, KRAS* remains challenging to target directly^20^. Therefore, disrupting downstream metabolic crosstalk mechanisms in PDA is a compelling alternative approach^21^.

In support of this idea, previous work from our lab described that PDA cells are uniquely dependent on KRAS*-mediated rewiring of glutamine metabolism for protection against oxidative stress^11^. Mitochondrial glutamate oxaloacetate transaminase 2 (GOT2) is critical for this rewired metabolism in PDA. In normal physiology, GOT2 functions in the malate-aspartate shuttle (MAS), a mechanism by which cells transfer reducing equivalents between the cytosol and mitochondria to balance the two independent NADH pools and maintain redox balance (**Fig.1A**). PDA cells driven by KRAS* divert metabolites from the MAS and increase flux through malic enzyme 1 (ME1) to produce NADPH^11^. Since this pathway is critical for PDA, we set out to evaluate GOT2 as a potential therapeutic target. This led to the observation that GOT2 was required for in vitro but not in vivo tumor growth. Ultimately, through metabolomics analyses and manipulation of the redox state in PDA cells, we discovered that pancreatic cancer-associated fibroblasts (CAFs) release pyruvate, which can be consumed by PDA cells to alleviate the redox imbalance induced by impaired mitochondrial metabolism. These data emphasize an under-appreciated role for GOT2 in pancreatic tumor redox homeostasis, and, perhaps more importantly, reinforce that the tumor microenvironment plays a major role in cancer metabolism and therapeutic resistance.

**Fig.1.**
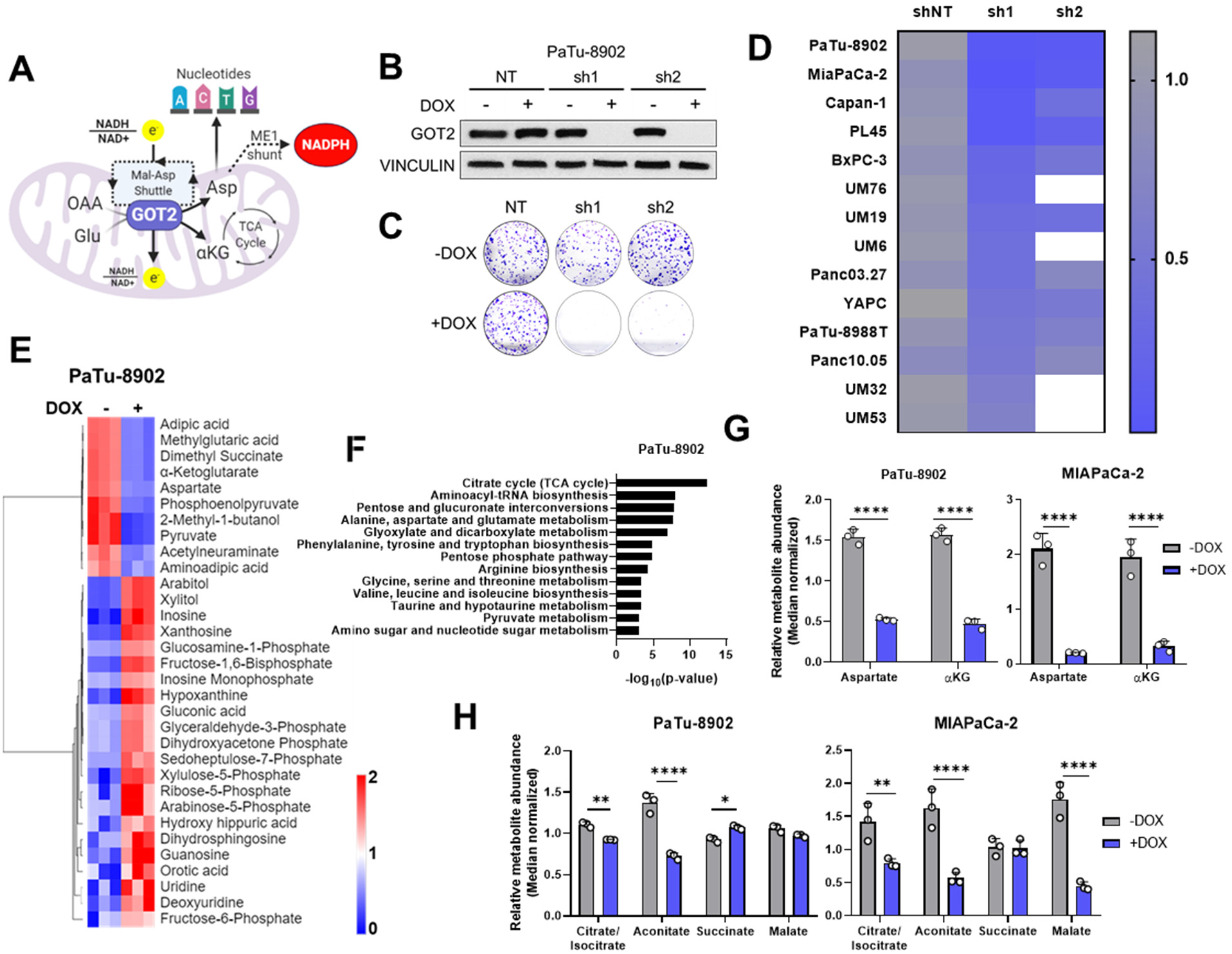
GOT2 KD impairs in vitro PDA colony formation. **A)** Schematic depicting the metabolic roles of GOT2 in PDA. **B)** Western blot of GOT2 expression with Vinculin loading control in PaTu-8902 iDox-shNT, shGOT2.1, or shGOT2.2 cells. **C)** Images of representative wells showing PaTu-8902 colony formation after GOT2 KD. **D)** Heatmap summarizing the relative colony formation after GOT2 KD across a panel of PDA cell lines, normalized to -DOX for each hairpin. **E-H)** Metabolites significantly changed between +DOX (n=3) and –DOX (n=3) (p<0.05, −1>log2FC>1) in PaTu-8902 iDox-shGOT2.1 cells as assessed by LC/MS. **E)** Heatmap depicting changes in relative metabolite abundances with 2D unsupervised hierarchical clustering. **F)** Metabolic pathways significantly changed as determined via Metaboanalyst. **G)** Relative abundances of Aspartate and αKG **H)** Relative abundances of TCA cycle intermediates. Bars represent mean ± SD, *p<0.05.

## RESULTS

### GOT2 is required for PDA colony formation in vitro

To expand on our previous work studying GOT2 in PDA^11^, and to evaluate GOT2 as a potential therapeutic target, we generated a panel of PDA cell lines with doxycycline-inducible expression of either a control non-targeting shRNA (shNT) or two independent shRNAs (sh1, sh2) targeting the GOT2 transcript. Cells cultured in media containing doxycycline (+DOX) exhibited a marked decrease in GOT2 protein expression compared to cells cultured in media without doxycycline (−DOX) (**Fig.1B; Extended Data Fig.1A**). This knockdown was specific for GOT2, relative to the cytosolic aspartate aminotransaminase GOT1 (**Extended Data Fig.1B**). Having validated GOT2 knockdown (KD), we tested the importance of GOT2 for cellular proliferation. In general, GOT2 KD in PDA cells impaired colony formation (**Fig.1C,D; Extended Data Fig.1C**). Consistent with our previous report^11^, GOT2 was not required for the proliferation of non-transformed pancreatic cell types (**Extended Data Fig.1D,E).**

Since GOT2 has several vital metabolic roles in a cell (**Fig.1A**), the changes caused by decreased GOT2 expression in PDA cells were examined using liquid chromatography coupled tandem mass spectroscopy (LC-MS/MS). Numerous changes in the intracellular metabolome of GOT2 KD cells were observed (**Fig.1E,F; Extended Data Fig.1F,G**). Of note, the products of the GOT2-catalyzed reaction, aspartate (Asp) and α-ketoglutarate (αKG), were decreased (**Fig.1G**). In addition to reduced αKG, there was a decrease in most TCA cycle intermediates, consistent with a role for GOT2 in facilitating glutamine anaplerosis (**Fig.1H**). These data demonstrate that loss of GOT2 in vitro depletes Asp and αKG, and also induces a strong growth inhibitory phenotype in vitro, prompting further exploration of GOT2 as a metabolic target in PDA in vivo.

### GOT2 is not required for PDA tumor growth in vivo

To test the effect of GOT2 KD on in vivo tumor growth, PDA cell lines were injected subcutaneously into the flanks of immunocompromised (NOD Scid gamma; NSG) mice. We allowed the tumors to establish for 7 days, after which the mice were fed normal chow or doxycycline chow *ad libitum*. Surprisingly, despite the inhibitory in vitro phenotype and robust suppression of GOT2 expression in vivo, tumors from five different cell lines grew unimpeded with GOT2 KD (**Fig.2A-C; Extended Data Fig.2A,B**). Nuclear Ki67 staining confirmed that tumors lacking GOT2 were proliferative, and actually displayed a modest, but significant, increase in Ki67-positive nuclei (**Fig.2D,E**). To further examine the role of GOT2 in a more physiologically relevant tumor model, PDA cells were injected orthotopically into the pancreas of NSG mice and tumors were allowed to establish for 7 days before feeding the mice regular or DOX chow. Similar to the subcutaneous model, GOT2 KD had no effect on the growth of orthotopic tumors (**Extended Data Fig.2C**).

**Fig.2.**
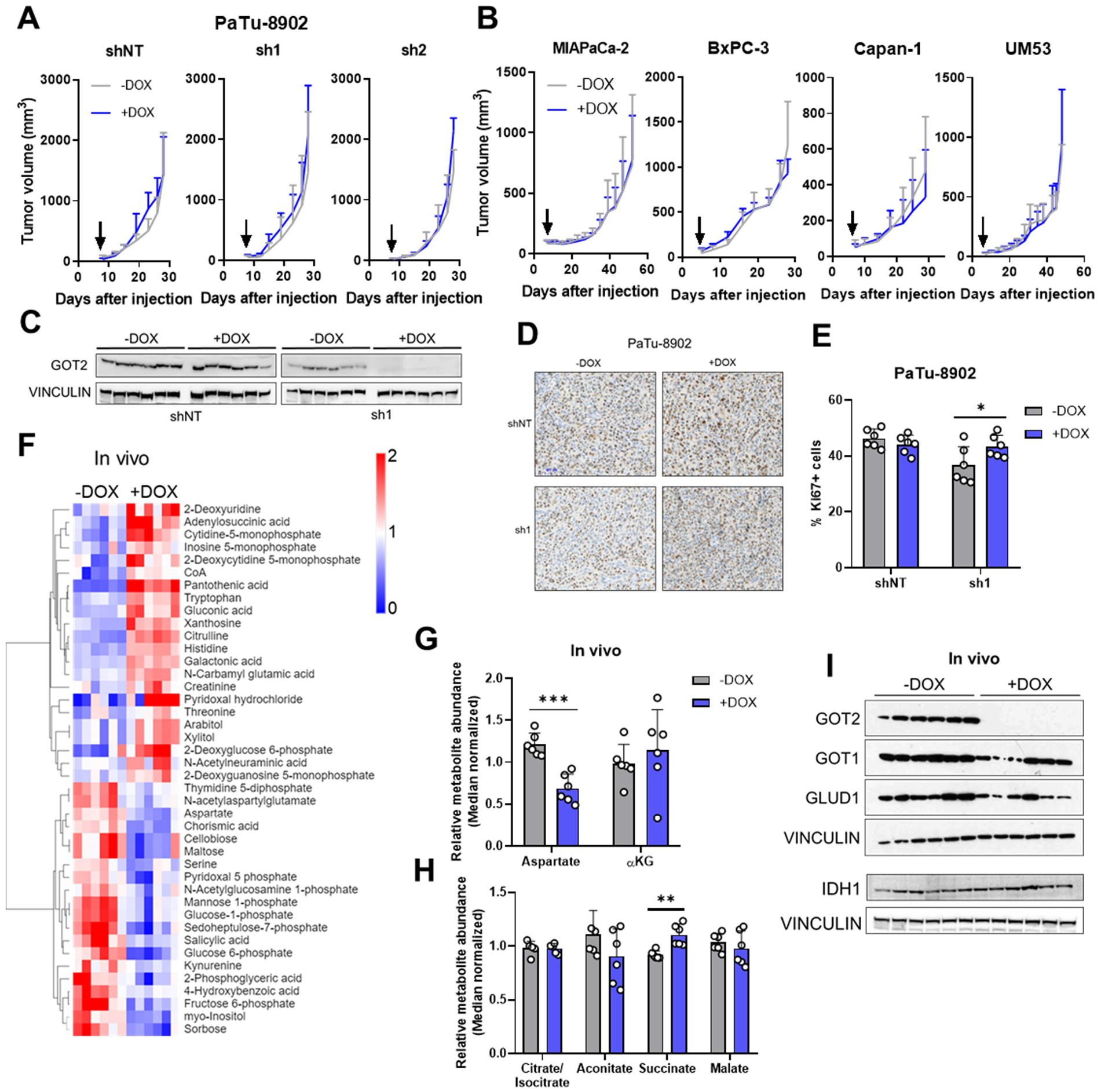
GOT2 is not required for PDA tumor growth in vivo. **A)** Growth of PaTu-8902 iDox-shNT, shGOT2.1, or shGOT2.2 subcutaneous xenografts in NSG mice (n=6 tumors per group). Arrows indicate administration of doxycycline chow. **B)** Growth across a panel of subcutaneous PDA iDox-shGOT2.1 subcutaneous xenografts in NSG mice (n=6 tumors per group). Arrows indicate administration of doxycycline chow. **C)** Western blot for expression of GOT2 and Vinculin loading control in PaTu-8902 iDox-shNT or shGOT2.1 subcutaneous xenografts. **D)** Representative images of Ki67 staining in tissue slices from PaTu-8902 iDox-shNT or shGOT2.1 subcutaneous xenografts, 10x. **E)** Quantification of nuclei positive for Ki67 in tissue slices from (B) (n=6 tumors per group). **F-H)** Metabolites significantly changed between PaTu-8902 iDox-shGOT2.1 +DOX (n=6) and –DOX (n=6) ((p<0.05, −0.5plog2FC>0.5) subcutaneous xenografts. **F)** Heatmap depicting changes in relative metabolite abundances with 2D unsupervised hierarchical clustering. **G)** Relative abundances of Aspartate and αKG. **H)** Relative abundances of TCA cycle intermediates. **I)** Western blots for expression of GOT1, GOT2, GLUD1, and IDH1, Vinculin loading control in PaTu-8902 iDox-shGOT2.1 tumors. Bars represent mean ± SD, *p<0.05, **p<0.01, ***p<0.001, ****p<0.0001.

Having observed a discrepancy between in vitro and in vivo dependence on GOT2 for proliferation, the relative abundances of intracellular metabolites from subcutaneous tumors were analyzed via LC-MS/MS to compare the metabolic changes between cell lines and tumors following loss of GOT2. While GOT2 KD induced some changes in tumor metabolite levels, the affected metabolic pathways were distinct from those observed in vitro, bearing in mind we were comparing homogenous cell lines with heterocellular xenografts (**Fig.2F; Extended Data Fig.2D**). Asp abundance was significantly decreased, yet αKG levels remained constant (**Fig.2G**), and TCA cycle intermediates were unaffected (**Fig.2H**). This led us to initially hypothesize that PDA cells rewire their metabolism in vivo to maintain αKG levels when GOT2 is knocked down. However, upon examination of the expression of other αKG-producing enzymes in GOT2 KD tumors, we did not observe a compensatory increase in expression (**Fig.2I**). Certainly, expression does not always dictate metabolic flux, but these data led us to adopt an alternative, cell-extrinsic hypothesis to explain the different in vitro and in vivo GOT2 KD phenotypes.

### Cancer-associated fibroblast conditioned media supports colony formation in GOT2 KD cells in vitro

Human PDA tumors develop a complex microenvironment composed of a tumor-promoting immune compartment, a robust fibrotic response consisting of diverse stromal cell types, and a dense extracellular matrix (ECM)^18^. The subcutaneous tumor milieu in immunocompromised mice is less complex than that of a human PDA tumor, α-smooth muscle actin (αSMA) staining revealed that activated fibroblasts comprised a substantial portion of the microenvironment in tumors regardless of GOT2 status (**Extended Data Fig.3A**). Additionally, we and others have previously reported mechanisms by which CAFs in the stroma engage in cooperative metabolic crosstalk with pancreatic cancer cells^22–24^. So, we hypothesized that CAFs were supporting PDA metabolism following GOT2 KD. To investigate potential metabolic crosstalk in a simplified setting, PDA cells were cultured in vitro with conditioned media (CM) from human CAFs (hCAFs). In support of our hypothesis, hCAF CM promoted colony formation in PDA cells with GOT2 KD in a dose-dependent manner (**Fig.3A,B; Extended Data Fig.3B**). Furthermore, hCAF CM displayed a more pronounced rescue phenotype compared to CM from tumor-educated macrophages (TEMs) or from PDA cells (**Extended Data Fig.3C**).

**Fig.3.**
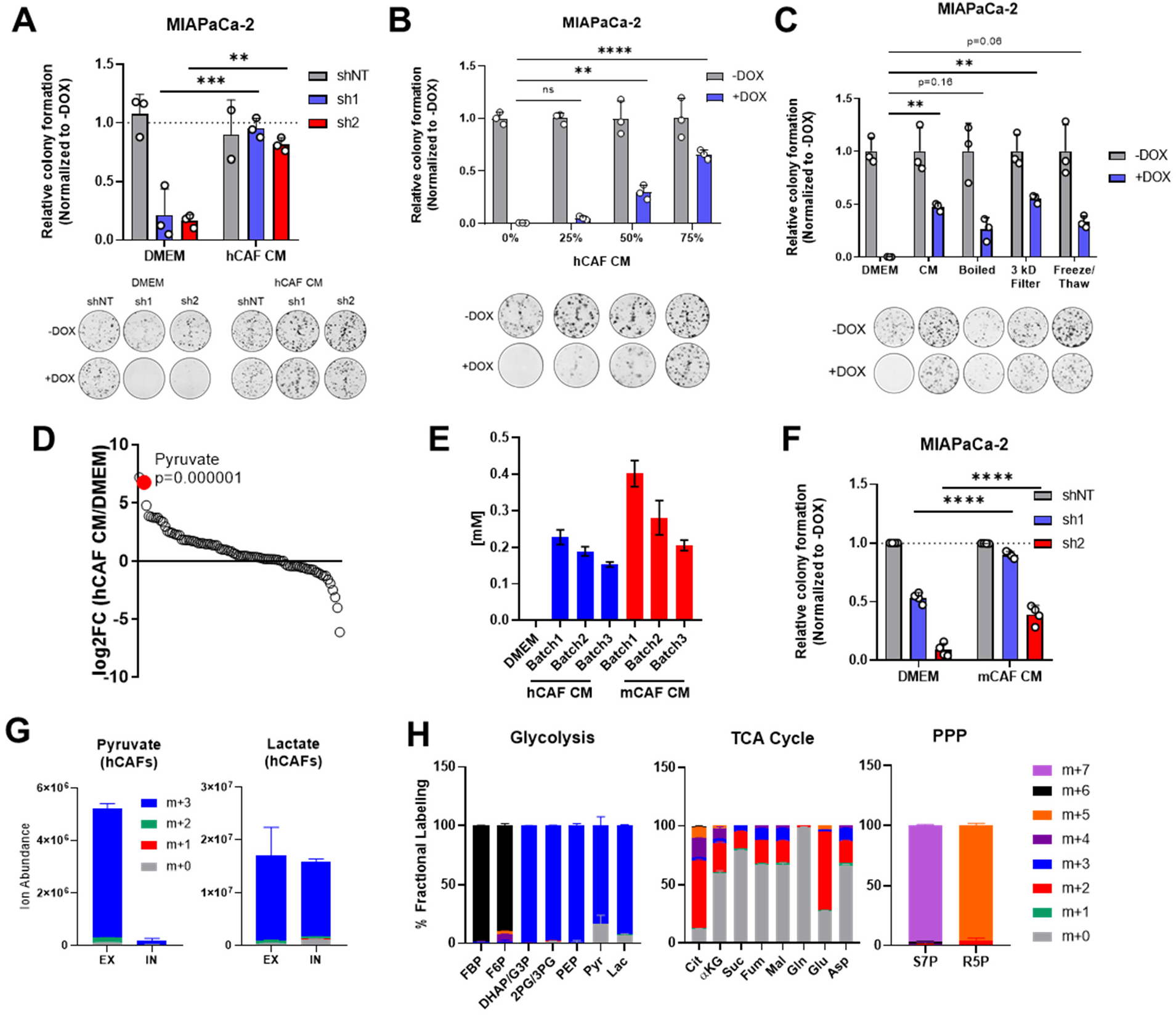
CAF CM restores PDA colony formation after GOT2 KD in vitro. **A)** Relative colony formation of MIAPaCa-2 iDox-shNT, shGOT2.1, or shGOT2.2 cells cultured in DMEM or hCAF CM, with images of representative wells. **B)** Relative colony formation of MIAPaCa-2 iDox-shGOT2.1 cells cultured in the indicated doses of hCAF CM, with images of representative wells. **C)** Relative colony formation of MIAPaCa-2 iDox-shGOT2.1 cells cultured in DMEM or fresh hCAF CM, boiled CM, CM passed through a 3 kD filter, or CM subjected to freeze/thaw cycles, with images of representative wells. **D)** Abundance of metabolites in hCAF CM compared to DMEM ranked according to log2FC. **E)** Quantification of pyruvate in three independently prepared hCAF or mCAF CM batches, as determined via LC/MS. **F)** Growth of MIAPaCa-2 iDox-shNT, shGOT2.1, or shGOT2.2 cells in DMEM or mCAF CM. **G)** Ion abundance of pyruvate and lactate isotopologues in intra and extracellular fractions from CAFs cultured in 13C-Glucose. **H)** Fractional labelling of intracellular isotopologues in glycolysis or TCA cyclefrom hCAFs cultured with U13C-Glucose. Fructose-6-phosphate (F6P), fructose-1,6-bisphosphate (FBP), dihydroxyacetone phosphate (DHAP), glyceraldehyde-3-phosphate (G3P), 2-phosphoglycerate (2PG), phosphoenol pyruvate (PEP), pyruvate (Pyr), lactate (Lac), citrate (Cit), succinate (Suc), fumarate (Fum), malate (Mal), glutamine (Gln), glutamate (Glu). sedoheptulose-7-phosphate (S7P), ribose-5-phosphate (R5P).Bars represent mean ± SD, **p<0.01, ***p<0.001, ****p<0.0001.

To begin to identify the factors in hCAF CM responsible for this effect, hCAF CM was boiled, filtered through a 3 kDa cut-off membrane, or subjected to cycles of freezing and thawing. In each of these conditions, hCAF CM supported colony formation in GOT2 KD cells, suggesting the relevant factor(s) was a metabolite (**Fig.3C; Extended Data Fig.3D**). The relative abundances of metabolites in hCAF CM determined via LC-MS/MS demonstrated that pyruvate was one of the most abundant metabolites released by hCAFs into the conditioned media, at a concentration near 250 μM (**Fig.3D,E; Extended Data Fig.3E**). Consistent with the idea of metabolite exchange, CAF-derived pyruvate was taken up by PDA cells, as cells cultured in hCAF CM had elevated levels of intracellular pyruvate (**Extended Data Fig.3F**).

In the in vivo model, mouse fibroblasts infiltrate the pancreatic xenografts and engage in crosstalk with cancer cells. Therefore, we tested whether our findings with hCAFs were also applicable in mouse cancer-associated fibroblasts (mCAFs). mCAFs isolated from a pancreatic subcutaneous xenograft in an NSG mouse also secreted pyruvate at ~250 μM in vitro (**Fig.3E**). Similar to hCAF CM, mCAF CM promoted PDA colony formation following GOT2 KD (**Fig.3F**, **Extended Data Fig.3G**). Since these mCAFs are the same CAFs encountered by PDA cells in our in vivo model, these data further support a mechanism by which CAFs compensate for loss of GOT2.

To better understand the production and release of pyruvate in CAFs, an isotope tracing experiment on CAFs with uniformly carbon labeled (U13C)-Glucose was performed. The pyruvate released by CAFs was indeed produced from glucose (**Fig.3G**), and, in support of previous studies^25,26^, these CAFs displayed labelling patterns indicative of glycolytic metabolism (**Fig.3H**).

### Pyruvate compensates for GOT2 KD in vitro

To determine whether pyruvate was the metabolite responsible for the rescue of GOT2 KD, cells were cultured in media supplemented with extracellular pyruvate. In a dose-dependent fashion, pyruvate increased colony formation in GOT2 KD cells (**Fig.4A,B**). This rescue was observed at both supra-physiological levels of pyruvate (1 mM) and at the levels we reported in CAF CM (250 μM). Furthermore, the concentration of pyruvate in mouse serum was previously reported to be 250 μM^27^, indicating that subcutaneous tumors are exposed to a dose of pyruvate that can rescue growth during GOT2 KD. Additionally, PDA cells expressing a genetically-encoded, fluorescent ATP sensor indicated that ATP levels dropped with GOT2 KD, and were restored with pyruvate supplementation (**Fig.4C**), reflecting the link between TCA cycle activity, respiration, and oxidative phosphorylation. The increase in ATP levels also correlated with an increase in overall proliferation (**Extended Data Fig.4A**). Furthermore, having identified a metabolite that permits in vitro proliferation of PDA cells without GOT2, we engineered CRISPR-Cas9 GOT2 knock out (KO) cells for further investigation (**Extended Data Fig.4B**). In support of the data generated using the doxycycline-inducible shRNA, GOT2 KO impaired colony formation of PDA cells, which was similarly restored through extracellular pyruvate supplementation (**Fig.4D; Extended Data Fig.4C**).

**Fig.4.**
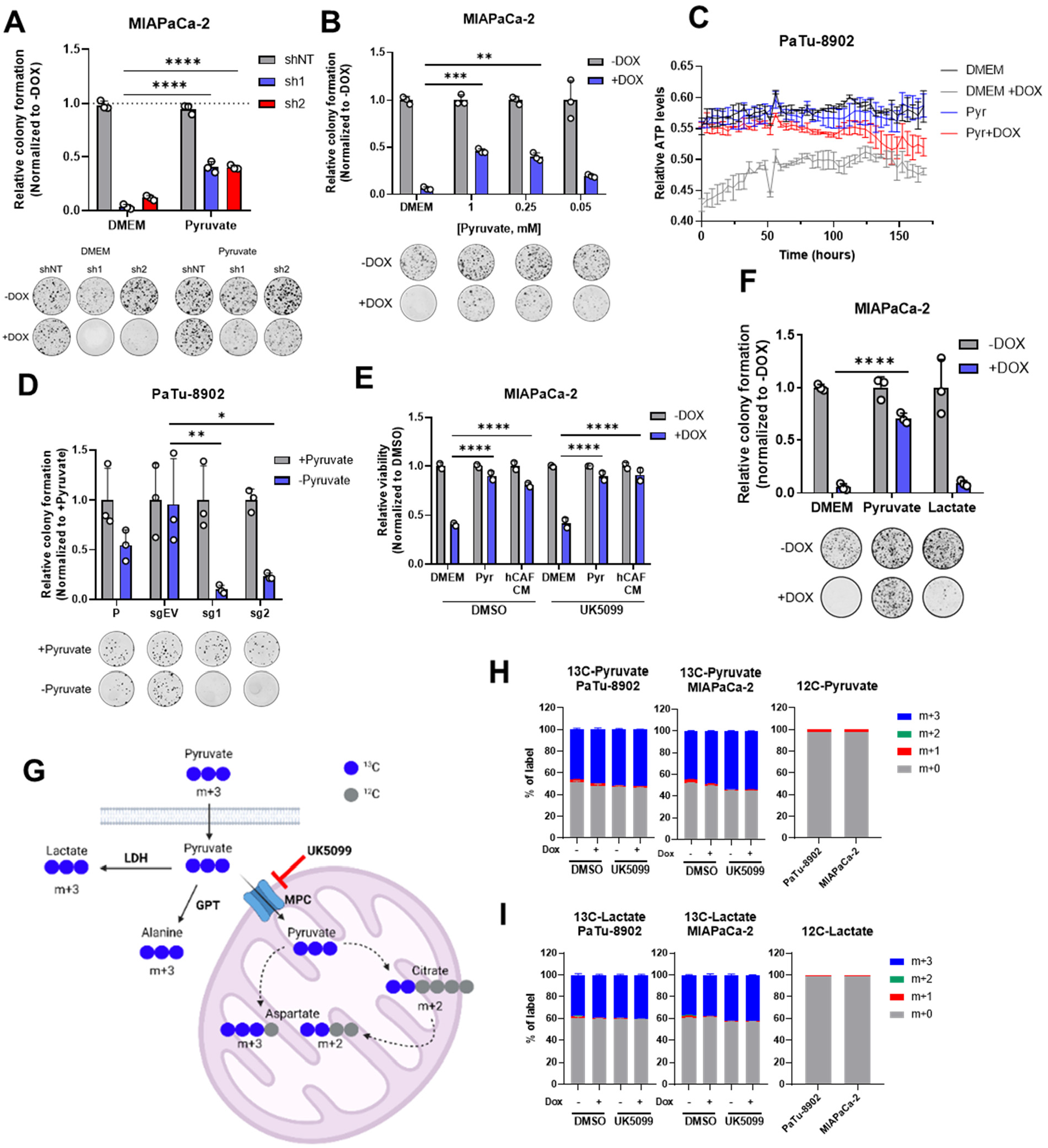
Pyruvate compensates for GOT2 KD in vitro. **A)** Relative colony formation of MIAPaCa-2 iDox-shNT, shGOT2.1, or shGOT2.2 cells cultured in DMEM or 1 mM pyruvate, with images of representative wells. **B)** Relative colony formation of MIAPaCa-2 iDox-shGOT2.1 cells cultured in DMEM or the indicated doses of pyruvate, with images of representative wells. **C)** Relative ATP levels over time in PaTu-8902 iDox-shGOT2.1 cells cultured in DMEM or 1 mM pyruvate. **D)** Relative colony formation of PaTu-8902 parental WT, sgEV, sgGOT2.1, or sgGOT2.2 cells cultured in DMEM or 1 mM pyruvate, with images of representative wells. **E)** Relative viability of MIAPaCa-2 iDox-shGOT2.1 cells after treatment with 5 μM UK5099 and culture in DMEM, 1 mM pyruvate, or hCAF CM. **F)** Relative colony formation of MIAPaCa-2 iDox-shGOT2.1 cells cultured in DMEM, 1 mM pyruvate, or 1 mM lactate. **G)** Schematic depicting the labelling patterns following incubation for 16 hours with 1 mM U13C-Pyruvate and treatment with 5 μM UK5099. Lactate dehydrogenase (LDH), mitochondrial pyruvate carrier (MPC), glutamate-pyruvate transaminase (GPT). **H,I)** Labelling patterns of intracellular pyruvate (H) or intracellular lactate (I) in PaTu-8902 or MIAPaCa-2 iDox-shGOT2.1 and treated with 5 μM UK5099. Bars represent mean ± SD, *p<0.05 **p<0.01, ***p<0.001, ****p<0.0001.

In addition to pyruvate, both Asp and αKG were present in hCAF CM (**Extended Data Fig.3E**). Previous studies have illustrated that Asp is rate limiting for proliferation^28–34^. Given that GOT2 is the predominant source of Asp in PDA cells (**Fig.1A**), we also tested if Asp, αKG, or the combination could rescue GOT2 KD. Although the combination afforded rescue, this required supraphysiological concentrations (4mM αKG, 10mM Asp). Interestingly, while rescue of GOT2 KD could be achieved with both Asp and αKG, pyruvate promoted GOT2 KD to an even greater degree than this combination (**Extended Data Fig.4D**). Considering that lack of rescue with single agent Asp could be explained by inefficient import by PDA cells, as has been reported^34^, we confirmed that PDA cells cultured with 10mM Asp displayed elevated levels of intracellular aspartate (**Extended Data Fig.4E,F**). Furthermore, providing cells with purine or pyrimidine nucleobases, metabolites downstream of Asp and responsible for proliferation (**Fig.1A**)^35^, did not restore colony formation after GOT2 KD (**Extended Data Fig.4G**). As such, these data demonstrate that while Asp and αKG can rescue GOT2 KD, several lines of evidence suggest that they are not responsible for the rescue imparted by hCAF CM.

Pyruvate is a pleiotropic molecule and has numerous fates and functions. These include roles in redox and nitrogen balance and as a fuel for myriad biosynthetic precursors and pathways^36^. Thus, we next set forth to determine the mechanism(s) by which pyruvate afforded rescue of GOT2 loss.

First, the potential for pyruvate to fuel mitochondrial metabolism was evaluated. As discussed above, GOT2 KD disrupted the TCA cycle, likely resulting from a decrease in αKG. Pyruvate can be converted to acetyl-CoA in the mitochondria by pyruvate dehydrogenase (PDH), where it enters the TCA cycle. Therefore, GOT2 KD cells were cultured with hCAF CM or pyruvate in the presence of the mitochondrial pyruvate carrier (MPC) inhibitor UK5099, which blocks entry of pyruvate into the mitochondria. Both hCAF CM and pyruvate retained the ability to rescue colony formation in cells with GOT2 KD in the presence of UK5099 (**Fig.4E**). Next, we examined lactate since it is found at millimolar concentrations in tumors and can be converted to pyruvate, as well as having the potential to be used as a carbon source for the TCA cycle^37^. Unlike pyruvate, lactate did not rescue colony formation following GOT2 KD (**Fig.4F**). Additionally, our group discovered previously that hCAFs release alanine, which is taken up by cancer cells and converted to pyruvate in the mitochondria by mitochondrial alanine aminotransaminase to fuel the TCA cycle^22^. Alanine similarly failed to rescue GOT2 KD (**Extended Data Fig.4H**). These data collectively indicated that even though GOT2 KD disrupts mitochondrial metabolism, pyruvate does not need to enter the mitochondria, and it appears that pyruvate is not directly participating in mitochondrial anaplerosis to promote colony formation.

To confirm our findings from these rescue experiments, we traced the metabolic fate of U13C-pyruvate in cells with GOT2 KD treated with UK5099 (**Fig.4G**). Media containing equal amounts of unlabelled glucose and labelled pyruvate was used to prevent washout of the pyruvate label by pyruvate generated via glycolysis from high glucose concentrations. This media formulation had no impact on the pyruvate GOT2 KD rescue phenotype (**Extended Data Fig.4I**).

In this experiment, roughly 50% of the intracellular pyruvate was labelled (**Fig.4H**), in line with the other half of the unlabelled pyruvate coming from glucose, and most of the labelled pyruvate was converted to lactate (**Fig.4I**). We did observe modest labelling of citrate, aspartate, and alanine from pyruvate (**Extended Data Fig.4J-L**). While the labelled TCA cycle and branching pathway intermediates decreased dramatically with UK5099, this MPC inhibition had no effect on the pyruvate GOT2 KD rescue phenotype. Together, the results from this experiment confirm that pyruvate does not need to enter the mitochondria and suggest a cytosolic rescue mechanism.

### GOT2 KD perturbs redox homeostasis in PDA cells

In cultured PDA cells, the MAS transfers glycolytic reducing potential to drive the electron transport chain (ETC) and maintain redox balance. We thus hypothesized that GOT2 KD interrupted this shuttle, preventing the proper transfer of electron potential in the form of NADH between these two compartments. Indeed, GOT2 KD increased the intracellular ratio of NADH to NAD+ (**Fig.5A**). Also, re-examination of the metabolomics dataset from cells with GOT2 KD revealed an impairment in glycolysis with a node at glyceraldehyde 3-phosphate dehydrogenase (GAPDH) (**Fig.5B**). GAPDH reduces NAD+ to produce NADH, where a build-up of NADH would serve to product-inhibit GAPDH activity. Indeed, this explains the metabolic signature observed, where metabolite pools in upstream glycolysis and branch pathways like the pentose phosphate pathway are increased, and those in downstream glycolysis are decreased (**Fig.5C**). On the contrary, this effect on glycolysis was not observed in the metabolomics analysis from subcutaneous GOT2 KD tumors, further illustrating the differential dependence on GOT2 in PDA in vitro and in vivo (**Extended Data Fig.5A**). The Seahorse glycolytic rate assay confirmed that glycolysis was indeed impaired during GOT2 KD in vitro (**Extended Data Fig.5B**).

**Fig.5.**
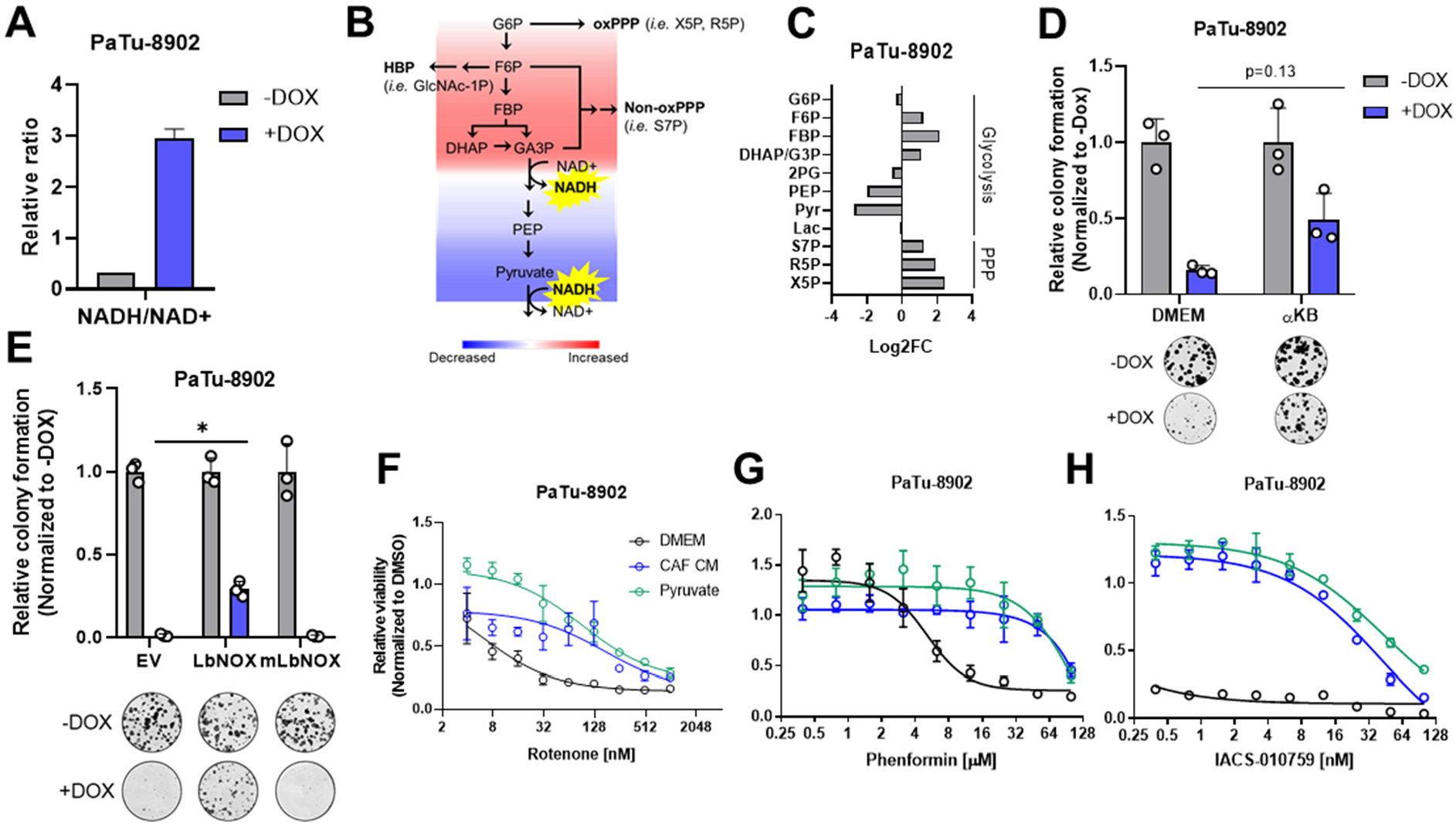
GOT2 KD disturbs redox homeostasis, which is restored by extracellular pyruvate. **A)** Relative ratio of NADH/NAD+ in PaTu-8902 iDox-shGOT2.1 cells, as determined by LC/MS. **B)** Schematic summarizing the metabolomics data following GOT2 KD depicting the effects of an increase in NADH levels on glycolysis. **C)** Log2 fold change of glycolytic and pentose phosphate pathway (PPP) intermediates in PaTu-8902 iDox-shGOT2.1 cells. **D)** Relative colony formation of PaTu-8902 iDox-shGOT2.1 cells cultured in DMEM or 1 mM αKB, with images of representative wells. **E)** Relative colony formation of PaTu-8902 iDox-shGOT2.1 cells expressing cytosolic or mitochondrial LbNOX. **F-H)** Relative viability of PaTu-8902 cells cultured in DMEM, hCAF CM, or 1 mM pyruvate and treated with the indicated doses of rotenone (F), phenformin (G), or IACS-010759 (H). Bars represent mean ± SD, *p<0.05Glucose-phosphate (DHAP), glyceraldehyde-3-phosphate (G3P), 2-phosphoglycerate (2PG), phosphoenol pyruvate (PEP), pyruvate (Pyr), lactate (Lac), sedoheptulose-7-phosphate (S7P), ribose-5-phosphate x(R5P), xylulose-5-phosphate (X5P).

### Pyruvate restores redox balance after GOT2 KD

NADH build-up leads to reductive stress, which can be relieved if the cell has access to electron acceptors^29^. Pyruvate is a notable metabolite in this regard, as it can accept electrons from NADH, producing lactate and regenerating NAD+ in a reaction catalyzed by lactate dehydrogenase (LDH). Since GOT2 KD caused a build-up of NADH, we hypothesized that this is the mechanism by which pyruvate rescues colony formation. To test this, GOT2 KD cells were cultured with α-ketobutyrate (αKB), another electron acceptor that turns over NADH in a mechanism analogous to pyruvate but without acting in downstream metabolism in the same fashion as pyruvate^31^. In support of our hypothesis, αKB also rescued colony formation after GOT2 KD (**Fig.5D; Extended Data Fig.5C**).

To test this further, GOT2 KD cells were generated to express either doxycycline-inducible cytosolic or mitochondrial Lactobacillus brevis NADH oxidase (LbNOX), which uses molecular oxygen to oxidize NADH and produce water and NAD+ (**Extended Data Fig.5D**)^38,39^. Cytosolic LbNOX, but not mitochondrial LbNOX, rescued colony formation imparted by GOT2 KD (**Fig.5E; Extended Data Fig.5E**). Curiously, the spatial control of this LbNOX system indicated that KD of mitochondrial GOT2 could be rescued by uniquely balancing the cytosolic NADH/NAD+ pool. Lastly, rescue of GOT2 KD requires the relief of NADH reductive stress, as opposed to increased production of NAD+, as supplementation with the NAD+ precursor nicotinamide mononucleotide (NMN) did not rescue GOT2 KD (**Extended Data Fig.5F**).

These findings provide clear evidence using several orthogonal strategies that GOT2 KD results in a build-up of NADH pools and reductive stress in PDA cells. Similarly, it has been well-established that inhibiting the activity of complex I of the ETC also results in an increase in NADH, which can be counteracted with extracellular pyruvate^29,31,34,39^. Therefore, since pyruvate is highly abundant in hCAF CM, we hypothesized that PDA cells cultured in hCAF CM would be protected from complex I inhibitors. Indeed, both hCAF CM and extracellular pyruvate conferred resistance to PDA cells against the complex I inhibitors rotenone, phenformin, and IACS-010759 (**Fig.5F-H**)^40^.

### Inhibiting pyruvate uptake and metabolism blocks rescue of GOT2 KD in vitro

According to our model, PDA cells are more vulnerable to GOT2 KD or complex I inhibitors in a pyruvate-depleted environment or if pyruvate uptake were blocked. Pyruvate can be transported by four MCT isoforms^41–43^, and an analysis of the CCLE database suggests that PDA cell lines primarily express MCT1 and MCT4 (**Extended Data Fig.6A**). Since MCT1 has a higher affinity for pyruvate than MCT4^42^, we decided to focus on MCT1 as the transporter by which PDA cells import pyruvate^44,45^. Indeed, PDA cells express significantly higher levels of MCT1, as compared to hCAFs (**Extended Data Fig.6B**). Similarly, examining expression of MCT1 from a recently published single cell analysis of a murine syngeneic orthotopic pancreatic tumor^46^ indicated that PDA cells express high levels of MCT1 (**Extended Data Fig.6C**).

Based on these expression data, we hypothesized that blocking pyruvate import through MCT1 would render cells vulnerable to GOT2 knockdown or complex I inhibition. The small molecule AZD3965 has specificity for MCT1 over MCT4^47–49^, therefore GOT2 KD cells were cultured in pyruvate or hCAF CM in the presence of AZD3965. In support of our hypothesis, neither pyruvate nor hCAF CM rescued GOT2 KD with MCT1 chemical inhibition (**Fig.6A,B**).

**Fig.6.**
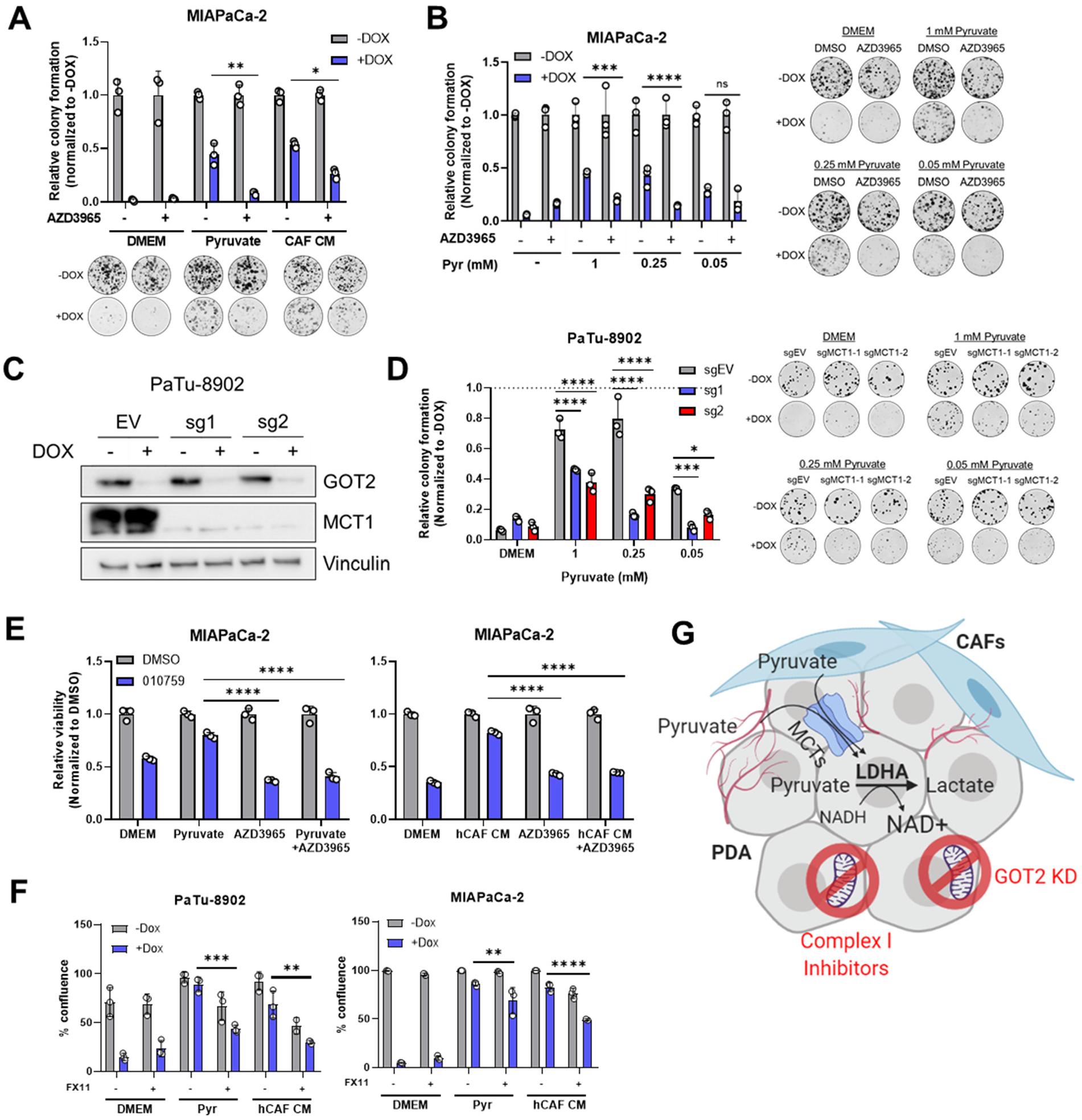
Blocking pyruvate uptake in vitro sensitizes PDA cells to redox disruption. **A)** Relative colony formation of MIAPaCa-2 iDox-shGOT2.1 cells treated with 100 nM AZD3965 and cultured in DMEM, 0.25 mM pyruvate, or hCAF CM, with images of representative wells. **B)** Relative colony formation of MIAPaCa-2 iDox-GOT2.1 cells treated with 100 nM AZD3965 and cultured in DMEM or the indicated doses of pyruvate, with images of representative wells. **C)** Western blot for GOT2 and MCT1, with Vinculin loading control, in PaTu-8902 iDox-shGOT2.1, MCT1 KO cells. **D)** Relative colony formation of PaTu-8902 iDox-shGOT2.1, MCT1 KO cells cultured in DMEM or with the indicated doses of pyruvate, with images or representative wells. **E)** Relative viability of MIAPaCa-2 cells treated with 100 nM IACS-010759 and 100 nM AZD3965 and cultured in DMEM, 1 mM pyruvate, or hCAF CM. **F)** Growth of PDA GOT2 KD cells cultured in 0.25 mM pyruvate or hCAF CM and treated with 25 μM FX11. **G)** Working model depicting the redox imbalance induced by GOT2 knockdown or complex I inhibition, which is then corrected through uptake of pyruvate released from CAFs or circulating through the vasculature. Bars represent mean ± SD, *p<0.05, **p<0.01, ***p<0.001, ****p<0.0001.

In parallel, MCT1 was knocked down in our DOX-inducible GOT2 KD cells (**Extended Data Fig.6D,F**). MCT1 KD modestly slowed the growth of GOT2 KD cells cultured in pyruvate or hCAF CM (**Extended Data Fig.6E,G**). We reasoned that partial knockdown could explain why MCT1 KD had a smaller effect than the chemical inhibitor MCT1. Therefore, we next generated MCT1 KO clones in our DOX-inducible GOT2 KD cells. Indeed, more complete disruption of MCT1 with CRISPR-Cas9 in GOT2 KD cells prevented pyruvate rescue (**Fig.6C,D**). We also demonstrate that MCT1 inhibition was most effective at physiological levels of pyruvate (250 μM) (**Fig.6B,D**). In parallel to experiments with GOT2 KD, MCT1 blockade was tested in combination with complex I inhibitors in PDA cells cultured in pyruvate or hCAF CM. AZD3965 also reversed the rescue activity of pyruvate or hCAF CM in PDA cells treated with IACS-010759 (**Fig.6E**).

The reduction of pyruvate to lactate by LDH is the central mechanism in our model by which NAD+ is regenerated. Thus, we next asked whether inhibiting LDH activity could also prevent GOT2 KD rescue by pyruvate or hCAF CM. Our PDA cell lines highly expressed the LDHA isoform of LDH, as determined by Western blotting (**Extended Data Fig.6H**). As such, we utilized the LDHA-specific chemical inhibitor FX11 in this study^50^. In further support of our model, inhibiting LDHA with FX11 slowed the in vitro growth of GOT2 KD cells cultured in pyruvate or hCAF CM, relative to single agent controls (**Fig.6F**, **Extended Data Fig.6I,J**).

Cumulatively, these data support an in vitro model whereby perturbation of mitochondrial metabolism with GOT2 KD or complex I inhibition disrupts redox balance in PDA cells. This can be restored through import of pyruvate from the extracellular environment and reduction to lactate to regenerate NAD+ (**Fig.6G**).

### Metabolic plasticity in vivo supports adaptation to combined GOT2 KD and inhibition of pyruvate metabolism

This proposed in vitro mechanism suggests that our initial observation, in which GOT2 is not required for in vivo tumor growth, could be explained by uptake of pyruvate from the TME by PDA cells. In vitro, MCT1 KD had a modest effect on the growth of GOT2 KD cells cultured in pyruvate or hCAF CM, yet MCT1 KD had no effect on the growth of GOT2 KD subcutaneous xenografts (**Extended Data Fig.7A-D**). Furthermore, MCT1 KO did not sensitize these tumors to GOT2 KD in vivo, as opposed to the results observed in vitro whereby MCT1 KO blocked the rescue of colony formation in GOT2 KD cells by pyruvate (**Extended Data Fig.7E,F**). Similarly, administration of AZD3965 did not impact the growth of GOT2 KD xenografts, again contrasting our in vitro data showing strong blockade of GOT2 KD rescue by pyruvate or hCAF CM (**Extended Data Fig.7G-I**). Lastly, LDHA inhibition in vitro blunted the GOT2 rescue activity of pyruvate and hCAF CM in vitro. However, administering FX11 to mice harboring GOT2 KD subcutaneous xenografts did not slow the growth of these tumors (**Extended Data Fig.8A-D**).

To evaluate the role of GOT2 in a more physiologically-relevant, immunocompetent model, we crossed the LSL-Kras^G12D^;Ptf1a-Cre (KC) mouse model with Got2^f/f^ mice to generate a LSL-Kras^G12D^;Got2^e^;Ptf1a-Cre mouse (KC-Got2^f/f^) (**Fig.7A, Extended Fig.9A**). We first confirmed that loss of *Got2* had no observable effect on the architecture or function in the healthy, non-transformed pancreas (**Extended Fig.9B-C**). This is in support of our data demonstrating loss of GOT2 was not deleterious in human, non-malignant pancreatic cell types.

**Fig.7.**
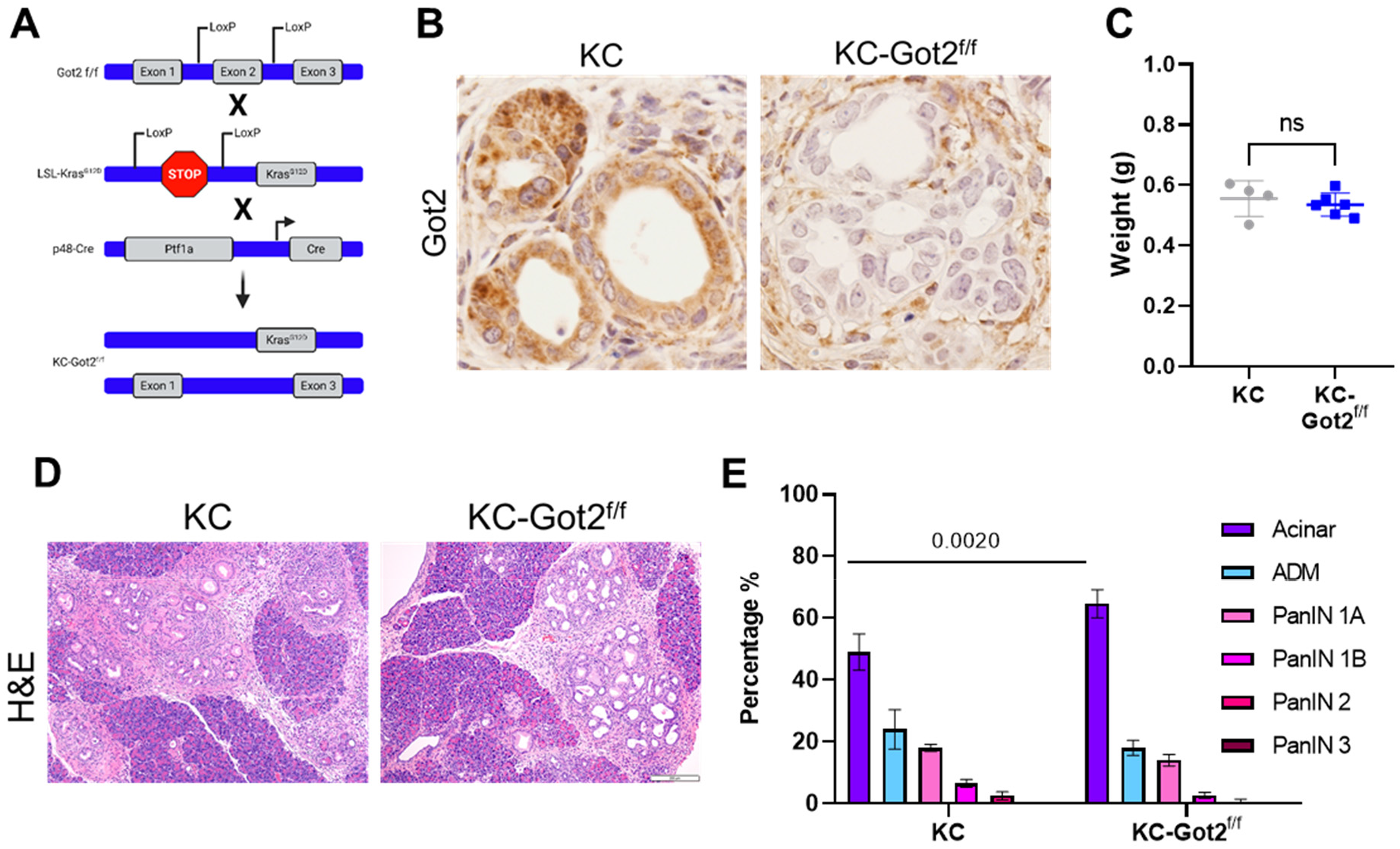
Got2 is not required for PDA tumorigenesis. **A)** Schematic depicting the breeding scheme for generating KC-Got2 mice. **B)** IHC for Got2 in pancreata from 3 month old KC or KC-Got2 mice, 40X. **C)** Weights of pancreata from 3 month old KC and KC-Got2 mice. **D)** Representative H&E images of pancreata from 3 month old KC or KC-Got2 mice, 10X. **E)** Scoring of pancreata from 3 month old KC (n=4) or KC-Got2 (n=6) mice. Bars represent mean ± SD.

KC-Got2 and KC controls were aged to 3 months, at which point pancreata were harvested. We confirmed loss of Got2 in the epithelial compartment via IHC (**Fig.7B**). No differences were observed in the weights of pancreata between 3 month KC-Got2 and KC mice (**Fig.7C**). Scoring the H&E stained tissues from these groups by a blinded pathologist revealed that KC-Got2 mice had a significantly greater percentage of healthy acinar cells compared to KC controls (**Fig.7D,E**). However, no differences were observed in the percentages of acinar-ductal metaplasia or PanIN grade between KC-Got2 and KC mice (**Fig.7E**). Additionally, we aged KC-Got2 mice to 6 months and compared the pancreata to matched 6 month KC historic controls. A histological analysis by a blinded pathologist did not identify a difference in number or severity of lesions (**Extended Data Fig.9D,E**). This suggests that loss of Got2 does not affect the progression of PDA following transformation by oncogenic Kras.

## DISCUSSION

GOT2 is an essential component of the MAS^51^ and required for redox homeostasis in PDA. Knockdown of GOT2 disrupts this shuttle and renders PDA cells incapable of transferring reducing equivalents between the cytosol and mitochondria, leading to a cytosolic build-up of NADH. This predominantly impacts the rate of glycolysis, an NAD+-coupled pathway, with secondary impacts on mitochondrial metabolism, that together slow the proliferation of PDA cells in vitro. For instance, GOT2 feeds into the ME1 shunt, which we demonstrated previously also produces pyruvate and sustains intracellular NADPH levels^52^. Extracellular supplementation with electron acceptors like pyruvate and αKB, or the expression of a cytosolic NADH oxidase, relieves NADH reductive stress and pathway feedback inhibition.

According to the data presented herein, GOT2 KD does not affect the growth of PDA tumors in vivo because electron acceptors in the tumor microenvironment can restore redox homeostasis. Indeed, pyruvate is present in mouse serum at 250 μM^27^, a concentration which is sufficient to compensate for GOT2 KD in vitro. Furthermore, we have shown that pancreatic CAFs release pyruvate, which is taken up and utilized by PDA cells. This is supported by previous findings in CAFs from other cancers^53^. Therefore, a source of pyruvate, either from CAFs or from the circulation, is available to PDA tumors.

This led us to hypothesize that blocking pyruvate uptake and metabolism would deprive PDA cells of a critical means to relieve the NADH stress mediated by GOT2 KD. In vitro, this hypothesis was supported when both inhibition of MCT1 and LDHA blocked the GOT2 KD rescue activity of pyruvate and CAF CM. However, this proved to be more complicated in vivo, as neither approach successfully sensitized tumors to GOT2 KD. We believe this could be explained by several mechanisms. First, while MCT4 has a lower affinity for pyruvate than MCT1, it can transport pyruvate^42^. It is also highly expressed in the cell lines used here (**Extended Data Fig.6A,C;7F**) and has been shown to confer resistance to cancer cells against MCT1 inhibition ^54,55^. Dual inhibition of MCT1 and MCT4 may be required to effectively block pyruvate uptake. Second, even with sufficient MCT blockade, PDA cells could still obtain pyruvate through other processes, such as macropinocytosis^7,9^. Third, while our study focuses on pyruvate, numerous circulating metabolites can function as electron acceptors and could relieve intracellular NADH build-up if cancer cells are unable to import pyruvate^56^. Fourth, reduction of pyruvate to lactate by LDH is not the only reaction by which NAD+ is regenerated. Recent studies have identified how serine pathway modulation, polyunsaturated fatty acid synthesis, and the glycerol-phosphate shunt all contribute to NADH turnover^57–59^. Future work remains to assess these mechanisms in PDA cells in vivo and if they compensate for an impaired MAS. Nevertheless, our data emphasize that redox homeostasis is a vital aspect of cancer cell metabolism and is maintained through a complex web of intracellular compensatory pathways and extracellular interactions.

Aside from its broader role in redox balance, GOT2 is also a prominent source of aspartate in PDA cells, and we demonstrate that GOT2 inhibition dramatically decreases aspartate levels both in vitro and in vivo. Previous studies have shown that aspartate availability is rate limiting in rapidly proliferating cells^30–32,34^. Surprisingly, even supraphysiological levels of aspartate as a single agent did not rescue GOT2 KD, and simultaneous treatment with supraphysiological doses of both Asp and αKG were required to provide a partial rescue of PDA cell proliferation in the absence of GOT2. Accordingly, this suggests that Asp does not participate in the ability to sustain tumor growth upon GOT2 KD in vivo, based on its availability at low micromolar concentrations and the limited uptake capacity of PDA cells^27^. Finally, supplementation of nucleosides similarly failed to rescue GOT2 KD. Therefore, we propose that the mechanism by which GOT2 KD slows PDA proliferation is through disruption of redox balance and deprivation of Asp. Supplying PDA cells with Asp meets a vital requirement for pyrimidine biosynthesis but does not address the redox imbalance. Pyruvate, on the other hand, regenerates NAD+ allowing broader metabolic processes to resume, including Asp production and nucleotide biosynthesis. In support of this, recent work in myoblasts demonstrated that while complex I inhibition with piericidin increased the NADH/NAD+ ratio leading to depletion of Asp, adding Asp back to the system neither restored redox balance nor induced proliferation^60^.

Our work also highlights the metabolic role of cancer-associated fibroblasts (CAFs) in PDA. Recent studies have shown that CAFs engage in cooperative metabolic crosstalk with cancer cells in many different tumor types^19,22,23,61,62^. We add to this body of literature by demonstrating that CAFs release pyruvate, which is taken up and utilized by PDA cells. However, much remains to be discovered about CAF metabolism. Some types of activated fibroblasts are known to be highly glycolytic^25,26^, an observation supported by our data. Yet the advent of single-cell RNA sequencing in murine pancreatic tumor models has led to a recent appreciation for the heterogeneity of CAFs^63–66^. The newly identified iCAF, myCAF, and apCAF populations have distinct functions in a pancreatic tumor^63^ and likely employ distinct metabolism to carry out these functions. Much more remains to be uncovered regarding competitive or cooperative interactions between PDA cells and the various CAF subpopulations, including which subtype(s) are responsible for pyruvate release.

Perhaps most importantly, this work emphasizes that the role of the tumor microenvironment must be considered when targeting cancer metabolism. Extracellular pyruvate serves to buffer redox imbalances induced by targeting mitochondrial metabolism. Indeed, approaches like disrupting the MAS shuttle via GOT2 KD or blocking complex I with small molecule inhibitors were less effective when PDA cells were cultured in supplemental pyruvate or in CAF CM. These data are relevant since numerous mitochondrial inhibitors are currently in clinical trials against solid tumor types (NCT03291938, NCT03026517, NCT03699319, NCT02071862). Previous studies have also shown that complex I inhibitors are more effective in combination with AZD3965^67^, a selective inhibitor of MCT1, though our work and others indicate that the status of other MCT isoforms should also be considered. Furthermore, the abundance of CAFs present in a tumor, as well as the level of circulating pyruvate in the patient, could predict outcomes for treatment with metabolic therapies that lead to redox imbalance. Targeting pancreatic cancer metabolism is an alluring approach, and a more detailed understanding of the metabolic crosstalk occurring in a pancreatic tumor can shed light on potential resistance mechanisms and inform more effective metabolic therapies.

## Acknowledgements

This work was funded by T32AI007413 and F31CA24745701 (SAK); CA148828 and CA245546 (YMS); a Pancreatic Cancer Action Network/AACR Pathway to Leadership award (13-70-25-LYSS), Junior Scholar Award from The V Foundation for Cancer Research (V2016-009), Kimmel Scholar Award from the Sidney Kimmel Foundation for Cancer Research (SKF-16-005), a 2017 AACR NextGen Grant for Transformative Cancer Research (17-20-01-LYSS), and NIH grants R37CA237421, R01CA248160, R01CA244931 (CAL).

Additional funding sources include F31CA254079 (JAJ); T32-DK094775 and T32-CA009676 (BSN); T32-GM11390 and F31-CA247076 (SBK); T32-CA009676, the American Cancer Society Postdoctoral Award PF-19-096-01, and the Michigan Institute for Clinical and Healthy Research (MICHR) Postdoctoral Translational Scholar Program fellowship award (NGS); the Michigan Postdoctoral Pioneer Program (ZCN); K99CA241357 and P30DK034933 (CJH); R01GM101171 (DBL); and Cancer Center support grant (P30 CA046592).

Metabolomics studies performed at the University of Michigan were supported by NIH grant DK097153, the Charles Woodson Research Fund, and the UM Pediatric Brain Tumor Initiative.

Research reported in this publication was supported by the National Cancer Institutes of Health under Award Number P30CA046592 by the use of the following Cancer Center Shared Resource(s): Flow Cytometry Core, Tissue and Molecular Pathology Core.

The University of Michigan Center for Gastrointestinal Research Core Director, Michael Mattea, provided assistance in tissue processing, sectioning, and staining.

## Author contributions

SAK, YMS, and CAL conceived of and designed this study. SAK, YMS, and CAL planned and guided the research and wrote the manuscript. SAK, LL, ALM, YZ, JAJ, PS, BC, AR, GT, BSN, SBK, NGS, MTH, HW, DL, SEA, JR, XG, LZ, AA, ZCN, SG, CJH, DBL, DRP-W, HY, HCC, MPdM, YMS, and CAL provided key reagents, performed experiments, analyzed, and interpreted data. YMS and CAL supervised the work carried out in this study.

## MATERIALS AND METHODS

### Cell culture

MIAPaCa-2, BxPC-3, Capan-1, Panc03.27, Panc10.05, PL45, and HPNE cell lines were obtained from ATCC. PaTu-8902, PaTu-8988T, and YAPC cells lines were obtained from DSMZ. UM6, UM19, UM28, UM32, UM53, and UM76 were generated from primary patient tumors at the University of Michigan. Human pancreatic stellate cells (hPSCs, also described here as hCAFs) were a generous gift from Rosa Hwang^68^. Mouse cancer-associated fibroblasts (mCAFs) were isolated as described below. All cell lines were cultured in high-glucose Dubelcco’s Modified Eagle Medium (DMEM, Gibco) without pyruvate and supplemented with 10% fetal bovine serum (FBS, Corning). 0.25% Trypsin (Gibco) was used to detach and passage cells. Cell lines were tested regularly for mycoplasma contamination using MycoAlert (Lonza). All cell lines in this study were validated for authentication using STR profiling via the University of Michigan Advanced Genomics Core. L-Aspartic acid (Sigma), dimethyl-α-ketoglutarate (Sigma), adenine (Sigma), guanine (Sigma), thymine (Sigma), cytosine (Sigma), sodium pyruvate (Invitrogen), α-ketobutyrate (Sigma), nicotinamide mononucleotide (NMN, Sigma), L-alanine (Sigma), and sodium lactate (Sigma) were used at the indicated concentrations. UK5099, AZD3965, and phenformin were purchased from Cayman chemical, Rotenone from Sigma, FX11 from MedChem Express, and IACS-010759 was a generous gift from Dr. Haoqiang Ying and Dr. Giulio Draetta, University of Texas MD Anderson Cancer Center.

### Doxy-inducible shGOT2 cells

The tet-pLKO-puro plasmid was a generous gift from Dmitri Wiederschain (Addgene #21915). Oligonucleotides encoding sense and antisense shRNAs (shGOT2.1-TRCN0000034824, shGOT2.2- TRCN0000034825) targeting GOT2 (NM_002080.4) were synthesized (Integrated DNA Technologies), annealed, and cloned at AgeI and EcoRI sites according to the Wiederschain Protocol^69^. A tet-pLKO non-targeting control vector (shNT- CCGGCAACAAGATGAAGAGCACCAACTCGAGTTGGTGCTCTTCATCTTGTTGTTTTT) was constructed using the same strategy. Tet-pLKO-shGOT2 and tet-pLKO-shNT lentiviruses were produced by the University of Michigan Vector Core using purified plasmid DNA. Stable cell lines were generated through transduction with optimized viral titers and selection with 1.5 μg/mL puromycin for 7 days.

### GOT2 or MCT1 knockout cells

GOT2 or MCT1 knockout PDA cell lines were generated using a CRISPR-Cas9 method described previously^70^. Briefly, sgRNA oligonucleotide pairs obtained from the Human GeCKO Library (v2, 3/9/2015). For GOT2 KO (sg1 (Fwd) 5’- CACCgAAGCTCACCTTGCGGACGCT-3’, (Rev) 5’-AAACAGCGTCCGCAAGGTGAGCTTc; sg2 (Fwd) 5’-CACCgCGTTCTGCCTAGCGTCCGCA-3’, (Rev) 5’- AAACTGCGGACGCTAGGCAGAACGc-3’), and for MCT1 KO (sg1 (Fwd) 5’- CACCgTGGGCCCGATTGGTCGCATG-3’, (Rev) 5’- AAACCATGCGACCAATCGGGCCCAc; sg2 (Fwd) 5’- CACCgTTTCTACAAGAGGCGACCAT-3’, (Rev) 5’- AAACATGGTCGCCTCTTGTAGAAAc-3’), were cloned into the pSpCas9(BB)-2A-Puro plasmid (PX459, v2.0; Addgene, #62988), transfected in PDA cell lines, and selected in puromycin for 7 days. Cells were then seeded into 96 well plates at a density of 1 cell per well, and individual clones were expanded. GOT2 knockout was verified via Western blot. Cells transfected with the empty PX459 vector were used as controls.

### MCT1 KD cells

Cells were transduced with 8 μg/mL polybrene and lentivirus containing the pGFP-C-shLenti plasmid (Origine, #TR30023) containing an shCTR sequence, shMCT1.1 [TL309405A (5′-GAGGAAGAGACCAGTATAGATGTTGCTGG-3′)], or shMCT1.2 [TL309405B (5′-ATCCAGCTCTGACCATGATTGGCAAGTAT-3′)]. These plasmids were a generous gift from Dr. Sean Morrison^49^. The cells were then centrifuged at 1000xg for 60 minutes at room temperature. Transduced cells were then expanded and sorted on the MoFlo Astrios (Beckman-Coulter). GFP+ cells were collected and expanded before verification of MCT1 KD via Western blotting.

### Transduction of LbNOX/mitoLbNOX

pINDUCER (Addgene, #44014) plasmids containing GFP, LbNOX, or mitoLbNOX were a generous gift from Dr. Haoqing Yang (MD Anderson). Plasmids were sequenced and transfected along with lentiviral packaging plasmids into HEK293FT cells with Lipofectamine 3000 (Thermo Fisher) per manufacturer’s instructions. Virus was collected after 48 hours and filtered through a 0.2 μm filter. PaTu-8902 and MIAPaCa-2 iDox-shGOT2.1 cells were seeded in 6 well plates at 250,000 cells/well, transduced with the indicated vectors, and selected in G418 at 500 μg/mL for 7 days. Expression of Flag-tagged LbNOX or mitoLbNOX was confirmed by Western blot with a Flag antibody after culturing cells in 1 μg/mL doxycycline for 3 days.

### Luciferase-expressing cells

MIAPaCa-2 iDox-shGOT2.1 cells were transduced with the FUGW-FL (EF1a-luc-UBC6-EGFP) lentiviral vector constructed previously^71^ and GFP+ cells were selected via flow cytometry. Luciferase activity was confirmed following transduction and selection with an in vitro luciferase assay and detection on a SpectraMax M3 Microplate reader (Molecular Devices).

### ATP fluorescent sensor/Incucyte growth assays

PaTu-8902 and MIAPaCa-2 iDox-GOT2.1 cells were transduced with CytoATP or CytoATP non-binding control vectors using the CytoATP Lentivirus Reagent Kit (Sartorious, #4772) and polybrene transfection reagent (Thermo Fisher) and selected for 7 days in 2 μg/mL puromycin. For proliferation and rescue experiments, cells were incubated in an Incucyte (Sartorious) equipped with a Metabolism Optical Module, where the ratio of ATP binding was detected and normalized to the non-binding control cells. Proliferation rate was determined by the percent confluence detected in the phase channel of the Incucyte normalized to Day 0 for each condition.

### Isolating mouse CAFs

UM2 subcutaneous xenografts from NSG mice were isolated and prepared in the laboratory of Dr. Diane Simeone, as reported previously ^72^, and single cell suspensions were plated and cultured *in vitro*. Mouse CAFs were separated from human pancreatic UM2 cancer cells using the Mouse Cell Depletion Kit (MACS Miltenyi Biotec) according to the manufacturer’s instructions.

### Conditioned media

Conditioned media was generated by splitting cells at ~90% confluence in a 10 cm^2^ plate into four 15 cm^2^ plates containing a final volume of 27 mL of growth media, and incubating for 72 hours at 37°C, 5% CO_2_. After, the media was collected in 50 mL conical tubes, centrifuged at 1,000 rpm for 5 minutes to remove any detached cells or debris, and divided into fresh 15 mL conical tubes in 10 mL aliquots before long-term storage at −80°C. For all conditioned media experiments, unless indicated otherwise, growth media was mixed with conditioned media for a final ratio of 75% conditioned media to 25% fresh growth media.

For the experiments in Fig.3 and Extended Data Fig.3, conditioned media were manipulated as follows. For boiling, the conditioned media tubes were placed in a water bath at 100°C for 10 minutes. To filter out factors >3 kDa, the conditioned media were transferred to a 3 kDa filter (Millipore) and centrifuged at 15,000 rpm in 30-minute increments until all the conditioned media had passed through the filter. To expose the conditioned media to freeze-thaw cycles, the tubes containing the conditioned media were thawed for 30 minutes in a 60°C water bath, and then frozen at −80°C for 30 minutes. This was repeated two more times for a total of three freeze-thaw cycles.

### Colony formation assays

Cells were seeded in 6 well plates at 200-400 cells per well in 2 mL of growth media and incubated overnight at 37°C, 5% CO_2_. The next day, the growth media was aspirated and fresh media containing the indicated compounds were added to the cells. Doxycycline was used at 1 μg/mL for all assays. For each assay, cells were incubated in the indicated conditions for 10 days, with the media and doxycycline changed every three days. After 10 days, the media was aspirated, the wells were washed once with PBS, and the cells were fixed in 100% methanol for 10 minutes. Next, the methanol was removed, and the cells were stained with 0.4% crystal violet for 10 minutes. Finally, the crystal violet was removed, the plates were washed under running water and dried on the benchtop overnight. The next day, images were taken of the plates with a Chemidoc BioRad imager and quantified using the ColonyArea plugin in ImageJ as described previously^73^.

### CyQUANT viability assay

Cells were seeded in 96 well black wall, clear bottom plates at 2,000 cells/well in 50 μL of media and incubated overnight at 37°C, 5% CO_2_. The next day, 150 μL of the indicated treatment media was added to the appropriate wells, and the cells were incubated for 4 more days. At endpoint, the media was removed from the wells, and the plates were stored overnight at −80°C. The next day, proliferation was determined by CyQUANT (Invitrogen) according to the manufacturer’s instructions, and fluorescence was detected on a SpectraMax M3 Microplate reader (Molecular Devices).

### Glycolytic rate assay

PaTu-8902 iDox-shGOT2.1 cells that had been cultured in 1 μg/mL doxycycline for 3 days were seeded at 2×10^4^ cells/well in 80 μl/well of normal growth media in an Agilent XF96 V3 PS Cell Culture Microplate (Agilent). To achieve an even distribution of cells within wells, plates were incubated on the bench top at room temperature for 1 hour before incubating at 37°C, 5% CO_2_ overnight. To hydrate the XF96 FluxPak (Agilent), 200 μL/well of sterile water was added and the entire cartridge was incubated at 37°C, CO_2_-free incubator overnight. The following day, one hour prior to running the assay, 60 μL of media was removed, and the cells were washed twice with 200 μL/well of assay medium (XF DMEM Base Medium, pH 7.4 containing 25 mM Glucose and 4 mM Glutamine; Agilent). After washing, 160 μL/well of assay medium was added to the cell culture plate for a final volume of 180 μL/well. Cells were then incubated at 37°C, in a CO_2_-free incubator until analysis. In parallel, one hour prior to the assay, water from the FluxPak hydration was exchanged for 200 μL/well of XF Calibrant 670 (Agilent) and the cartridge was returned to 37°C, CO_2_-free incubator until analysis. Rotenone/Antimycin (50 μM, Agilent) and 2DG (500 mM, Agilent) were re-constituted in assay medium to make the indicated stock concentrations. 20 μL of rotenone/antimycin was loaded into Port A for each well of the FluxPak and 22 μL of 2DG into Port B, for a final concentration of 0.5 μM and 50 mM, respectively. The Glycolytic Rate Assay was conducted on an XF96 Extracellular Flux Analyzer (Agilent) and PER was calculated using Wave 2.6 software (Agilent). Following the assay, PER was normalized to cell number with the CyQUANT NF Cell Proliferation Assay (Invitrogen) according to manufacturer’s instructions.

### Protein lysates

Cell lines cultured in 6 well plates in vitro were washed with ice-cold PBS on ice and incubated in 250 μL of RIPA buffer (Sigma) containing protease (Roche) and phosphatase (Sigma) inhibitors on ice for 10 minutes. Next, cells were scraped with a pipet tip, and the resulting lysate was transferred to a 1.5 mL tube also on ice. The lysate was centrifuged at 15,000 rpm for 10 minutes at 4°C. After, the supernatant was transferred to a fresh 1.5 mL tube and stored at −80°C.

In vivo tumor tissue was placed in a 1.5 mL tube containing a metal ball and 300 μL RIPA buffer with protease and phosphatase inhibitors. The tissue was homogenized using a tissue lyser machine. Then, the resulting lysate was centrifuged at 15,000 rpm for 10 minutes at 4°C. After, the supernatant was transferred to a fresh 1.5 mL tube and stored at −80°C.

### Western blotting

Protein levels were determined using a BCA assay (Thermo Fisher), according to manufacturer’s instructions. Following quantification, the necessary volume of lysate containing 30 μg of protein was loading dye (Invitrogen) and reducing agent (Invitrogen) and incubated at 90°C for 5 minutes. Next, the lysate separated on a 4-12% Bis-Tris gradient gel (Invitrogen) along with a protein ladder (Invitrogen) at 150 V until the dye reached the bottom of the gel (about 90 minutes). Then, the protein was transferred to a methanol-activated PVDF membrane (Millipore) at 25 V for 1 hour. After that, the membrane was blocked in 5% blocking reagent (Biorad) dissolved in TBS-T rocking for > 1 hour. Next, the membrane was incubated overnight at 4°C rocking in the indicated primary antibody diluted in blocking buffer. The next day, the primary antibody was removed, and the membrane was washed 3 times in TBS-T rocking for 5 minutes. Then, the membrane was incubated for 1 hour rocking at room temperature in the appropriate secondary antibody diluted in TBS-T. Finally, the membrane was washed as before, and incubated in Clarity ECL reagent (Biorad) according to manufacturer’s instructions before imaging on a Biorad Chemidoc. The following primary antibodies were used in this study: GOT2 (Atlas, HPA018139), GOT1 (Abcam, ab171939), GLUD1 (Abcam, ab166618), IDH1 (Cell Signaling, 3997S), MCT1 (Abcam, ab85021), MCT4 (Sigma, AB3316P), anti-Flag (Sigma, F3165), Vinculin (Cell Signaling, 13901S), LDHA (Cell Signaling, 3582), LDHB (Abcam, ab53292), and the anti-rabbit-HRP secondary antibody (Cell Signaling, 7074S)

### Isolating polar metabolites

For intracellular metabolome analyses, cells were seeded at 10,000 cells in 2 mL of growth media per well of a 6 well plate and incubated overnight. The next day, the growth media was removed, and cells were incubated in media containing the indicated compounds for 6 days, with the media being changed every 3 days. On day 6, the media was removed, and the cells were incubated in 1 mL/well of ice-cold 80% methanol on dry ice for 10 minutes. Following the incubation, the wells were scraped with a pipet tip and transferred to a 1.5 mL tube on dry ice.

To analyze extracellular metabolomes, 0.8 mL of ice-cold 100% methanol was added to 0.2 mL of media, mixed well, and incubated on dry ice for 10 minutes.

The tubes were then centrifuged at 15,000 rpm for 10 minutes at 4°C, and the resulting metabolite supernatant was transferred to a fresh 1.5 mL tube. The metabolites were then dried on a SpeedVac until all the methanol had evaporated, and re-suspended in a 50:50 mixture of methanol and water.

### Snapshot metabolomics

Samples were run on an Agilent 1290 Infinity II LC −6470 Triple Quadrupole (QqQ) tandem mass spectrometer (MS/MS) system with the following parameters: Agilent Technologies Triple Quad 6470 LC-MS/MS system consists of the 1290 Infinity II LC Flexible Pump (Quaternary Pump), the 1290 Infinity II Multisampler, the 1290 Infinity II Multicolumn Thermostat with 6 port valve and the 6470 triple quad mass spectrometer. Agilent Masshunter Workstation Software LC/MS Data Acquisition for 6400 Series Triple Quadrupole MS with Version B.08.02 is used for compound optimization, calibration, and data acquisition.

Solvent A is 97% water and 3% methanol 15 mM acetic acid and 10 mM tributylamine at pH of 5. Solvent C is 15 mM acetic acid and 10 mM tributylamine in methanol. Washing Solvent D is acetonitrile. LC system seal washing solvent 90% water and 10% isopropanol, needle wash solvent 75% methanol, 25% water. GC-grade Tributylamine 99% (ACROS ORGANICS), LC/MS grade acetic acid Optima (Fisher Chemical), InfinityLab Deactivator additive, ESI –L Low concentration Tuning mix (Agilent Technologies), LC-MS grade solvents of water, and acetonitrile, methanol (Millipore), isopropanol (Fisher Chemical).

An Agilent ZORBAX RRHD Extend-C18, 2.1 × 150 mm and a 1.8 um and ZORBAX Extend Fast Guards for UHPLC are used in the separation. LC gradient profile is: at 0.25 ml/min, 0-2.5 min, 100% A; 7.5 min, 80% A and 20% C; 13 min 55% A and 45% C; 20 min, 1% A and 99% C; 24 min, 1% A and 99% C; 24.05 min, 1% A and 99% D; 27 min, 1% A and 99% D; at 0.8 ml/min, 27.5-31.35 min, 1% A and 99% D; at 0.6 ml/min, 31.50 min, 1% A and 99% D; at 0.4 ml/min, 32.25-39.9 min, 100% A; at 0.25 ml/min, 40 min, 100% A. Column temp is kept at 35 °C, samples are at 4 °C, injection volume is 2 μl.

6470 Triple Quad MS is calibrated with the Agilent ESI-L Low concentration Tuning mix. Source parameters: Gas temp 150 °C, Gas flow 10 l/min, Nebulizer 45 psi, Sheath gas temp 325 °C, Sheath gas flow 12 l/min, Capillary −2000 V, Delta EMV −200 V. Dynamic MRM scan type is used with 0.07 min peak width, acquisition time is 24 min. dMRM transitions and other parameters for each compounds are list in a separate sheets. Delta retention time of plus and minus 1 min, fragmentor of 40 eV and cell accelerator of 5 eV are incorporated in the method.

The MassHunter Metabolomics Dynamic MRM Database and Method was used for target identification. Key parameters of AJS ESI were: Gas Temp: 150 °C, Gas Flow 13 l/min, Nebulizer 45 psi, Sheath Gas Temp 325 °C, Sheath Gas Flow 12 l/min, Capillary 2000 V, Nozzle 500 V. Detector Delta EMV(−) 200.

The QqQ data were pre-processed with Agilent MassHunter Workstation QqQ Quantitative Analysis Software (B0700). Each metabolite abundance level in each sample was divided by the median of all abundance levels across all samples for proper comparisons, statistical analyses, and visualizations among metabolites. The statistical significance test was done by a two-tailed t-test with a significance threshold level of 0.05.

Heatmaps were generated and data clustered using Morpheus Matrix Visualization and analysis tool (https://software.broadinstitute.org/morpheus).

Pathway analyses were conducted using MetaboAnalyst (https://www.metaboanalyst.ca).

### U13C-Glucose, U13C-Pyruvate isotope tracing

For glucose tracing, CAFs seeded were seeded in 6 well plates at 2×10^5^ cells/well and incubated for 72 hours in growth media containing U13C-Glucose (Cambridge Isotope Laboratories).

For pyruvate tracing, PaTu-8902 and MIAPaCa-2 iDox-shGOT2.1 cells were cultured in media containing 1 mM unlabelled glucose and 1 mM U13C-Pyruvate (Cambridge Isotope Laboratories) for 16 hours.

Polar metabolites were extracted from the media and cells according to the method described above. Isotope tracing experiments utilized the same chromatography as described in the Snapshot Metabolomics section, and were conducted on two instruments with the following parameters:

Agilent Technologies Q-TOF 6530 LC/MS system consists of a 1290 Infinity II LC Flexible Pump (Quaternary Pump), 1290 Infinity II Multisampler, 1290 Infinity II Multicolumn Thermostat with 6 port valve and a 6530 Q-TOF mass spectrometer with a dual Assisted Jet Stream (AJI) ESI source. Agilent MassHunter Workstation Software LC/MS Data Acquisition for 6200 series TOF/6500 series Q-TOF Version B.09.00 Build 9.0.9044.a SP1 is used for calibration and data acquisition.

An Agilent ZORBAX RRHD Extend-C18, 2.1 × 150 mm and a 1.8 um and ZORBAX Extend Fast Guards for UHPLC are used in the separation. LC gradient profile is: at 0.25 ml/min, 0-2.0 min, 100% A; 12.00 min, 1% A and 99% C; 16.00 min 1% A and 99% C; 18.00 min, 1% A and 99% D; 19.30 min, 1% A and 99% C; 19.90 min, 1% A and 99% D (0.80 ml/min); 22.45 min, 1% A and 99% D (0.8 ml/min); 22.65 min, 1% A and 99% D (0.4 ml/min); 29.35 min, 1% A and 99% D (0.4 ml/min); 29.45 min, 100% A (0.25 ml/min), 40 min. Column temp is kept at 35 C, samples are at 4 °C, injection volume is 5 μl.

Solvent A is 97% water and 3% methanol 15 mM acetic acid and 10 mM tributylamine at pH of 5. Solvent C is 15 mM acetic acid and 10 mM tributylamine in methanol. Washing Solvent D is acetonitrile. LC system seal washing solvent 90% water and 10% isopropanol, needle wash solvent 75% methanol, 25% water.

Agilent 6530 Q-TOF MS is calibrated with ESI-L Low Concentration Tuning mix. Source parameters: Gas temp 250 °C, Gas flow 13 l/min, Nebulizer 35 psi, Sheath gas temp 325 °C, Sheath gas flow 12 l/min, Vcap 3500 V, Nozzle Voltage (V) 1500, Fragmentor 140, Skimmer1 65, OctopoleRFPeak 750. The MS acquisition mode is set in MS1 with mass range between 50-1200 da with collision energy of zero. The scan rate (spectra/sec) is set at 1 Hz. The LC-MS acquisition time is 18 min and total run time is 30 min. Reference masses are enabled with reference masses in negative mode of 112.9856 and 1033.9881 da.

Agilent Technologies 6545B Accurate-Mass Quadrupole Time of Flight (MS Q-TOF) LC/MS coupled with an Agilent 1290 Infinity II UHPLC. Agilent Masshunter Workstation Software LC/MS Data Acquisition for 6500 Series QTOF MS with Version 09.00, Build9.0.9044.0 was used for tuning, calibration, and data acquisition.

In negative mode, the UHPLC was configured with 1290 Infinity II LC Flexible Pump (Quaternary Pump),1290 Infinity II Multisampler, 1290 Infinity II Multicolumn Thermostat with 6 port valves. Solvent Ais97% water and 3% methanol 15 mM acetic acid and 10 mM tributylamine at pH of 5. Solvent C is 15 mMacetic acid and 10 mM tributylamine in methanol. Washing Solvent D is acetonitrile. LC system seal washing solvent 90% water and 10% isopropanol, needle wash solvent 75% methanol, 25% water. GC-grade Tributylamine 99% (ACROS ORGANICS), LC/MS grade acetic acid Optima (Fisher Chemical), InfinityLab Deactivator additive, ESI–L Low concentration Tuning mix (Agilent Technologies), LC-MS grade solvents of water, and acetonitrile, methanol (Millipore), isopropanol (Fisher Chemical).

Agilent ZORBAX RRHD Extend-C18, 2.1 × 150 mm, 1.8 um and ZORBAX Extend Fast Guards for UHPLC are used in the separation. LC gradient profile is: at 0.25 ml/min, 0-2.5 min, 100% A; 7.5min, 80% A and 20% C; 13min 55% A and 45% C; 20 min, 1% A and 99% C; 24 min, 1% A and 99% C; 24.05 min, 1% A and 99% D; 27min, 1% A and 99% D; at 0.8 ml/min, 27.5-31.35 min, 1% A and 99% D; at 0.6 ml/min, 31.50 min, 1% A and 99% D; at 0.4 ml/min, 32.25-39.9 min, 100% A; at 0.25 ml/min, 40 min, 100% A. Column temp is kept at 35°C, samples are at 4°C, injection volume is 5 μl. In negative scan mode, the Agilent G6545B Q-TOF MS with Dual AJD ESI Sources in centroid mode was configured with following parameters: Acquisition range: 50-1200 da at scan rate of 1 spectra/sec, Gas temp 250 °C, Gas Flow 13 L/min, Nebulizer at 40 psi, Sheath Gas Heater 325 °C, Sheath Gas Flow 12L/min, Capillary 3500 V, Nozzle Voltage 1000 V, Fragmentor 130 V, Skimmer1 60 V, Octopole RFPeak 750V, Collision 0 V, Auto Recalibration limit of detection 150 ppm with min height 1000 counts, Reference ions of two at 59.0139 and 980.0164 da.

Data processing was performed in Agilent MassHunter Workstation Profinder 10.0 Build 10.0.10062.0. Isotopologue distributions were derived from a compound standard library built in Agilent MassHunter PCDL (Personal Compound and Database Library) v7.0.

### Xenograft studies

Animal experiments were conducted in accordance with the Office of Laboratory Animal Welfare and approved by the Institutional Animal Care and Use Committees of the University of Michigan. NOD scid gamma (NSG) mice (Jackson Laboratory) 6-10 weeks old of both sexes were maintained in the facilities of the Unit for Laboratory Animal Medicine (ULAM) under specific pathogen-free conditions.

Cells expressing doxycycline-inducible shNT or doxycycline-inducible shGOT2 were injected subcutaneously into both the left and right flanks of male and female NSG mice, with 1-4×10^6^ cells in a mixture of 50 μL media and 50 μL Matrigel (Corning) per injection. Tumors were established for 7 days before mice were fed either normal chow or chow containing doxycycline (BioServ). Tumors were measured with calipers two times per week, and mice were euthanized once the tumors reached a diameter of 2 cm^3^. Subcutaneous tumor volume (V) was calculated as V=1/2(length × width^2^). At endpoint, the tumors were removed, and fragments were either snap frozen in liquid nitrogen and stored at −80°C or fixed in ZFix solution (Anatech) for histology.

Cells expressing luciferase in addition to doxycycline-inducible shGOT2 were injected into the pancreas tail of NSG mice, with 200,000 cells in a mixture of 50 μL media and 50 μL Matrigel (Corning) per injection. Tumors were established for 7 days before mice were fed either normal chow or chow containing doxycycline (BioServ). Tumor progression was monitored by weekly intraperitoneal injections of luciferin (Promega) and bioluminescence imaging (BLI) on an IVIS SpectrumCT (Perkin Elmer). BLI was analyzed with Living Image software (PerkinElmer) At endpoint, the tumors were removed, and fragments were either snap frozen in liquid nitrogen and stored at −80°C or fixed in ZFix (Anatech) solution for histology.

AZD3965 or FX11 was dissolved in DMSO and stored at −80’C in aliquots. Each day, one aliquot was thaw and mixed with a 0.5% Hypromellose (Sigma), 0.2% tween80 (Sigma) solution such that the final DMSO concentration was 5%. Vehicle or AZD3965 was administered at 100 mg/kg by daily oral gavage. FX11 was administered at 2 mg/kg by daily intraperitoneal injection.

### Histology

Tissues were processed using a Leica ASP300S tissue processor (Leica Microsystems). Paraffin-embedded tissues were sectioned at 4 μm and stained for specific target proteins using the Discovery Ultra XT autostainer (Ventana Medical Systems), with listed antibodies, and counterstained with Mayer’s hematoxylin (Sigma). Hematoxylin and eosin (H&E) staining was performed using Mayer’s hematoxylin solution and Eosin Y (Thermo Fisher). IHC slides were then scanned on a Pannoramic SCAN scanner (Perkin Elmer). Scanned images were quantified using algorithms provided from Halo software version 2.0 (Indica Labs). The following antibodies were used for IHC: Ki67 (1:1,000; Abcam, ab15580), αSMA (1:20,000; Abcam, ab5694), Got2 (1:500; Atlas, HPA018139).

### KC-Got2^f/f^ model

Mice containing loxP sites flanking exon 2 of the Got2 gene were generated by Ozgene. These mice were crossed to the LSL-Kras^G12D^;Ptf1a-Cre model^74^. Tails from 3-week-old mice were collected at weaning and submitted to Transnetyx for genotyping. The following primers were used: Kras^G12D^ (Fw-GGCCTGCTGAAAATGACTGAGTATA, Rev- CTGTATCGTCAAGGCGCTCTT); Got2 WT (Fw- GCAGATTAAAACCACAAGGCCTGTA, Rev- ATGTTAAAATTGTCATCCCCTTGTGC); Got2 floxed (Fw- GCAGATTAAAACCACAAGGCCTGTA,Rev-AGAGAATAGGAACTTCGGAATAGGAACT); Cre (Fw-TTAATCCATATTGGCAGAACGAAAACG, Rev-CAGGCTAAGTGCCTTCTCTACA).

### Statistics

Statistics were performed using Graph Pad Prism 8. Groups of 2 were analyzed with two-tailed students t test, groups greater than 2 were compared using one-way ANOVA analysis with Tukey post hoc test or two-way ANOVA with Dunnett’s correction for multiple independent variables. All error bars represent mean with standard deviation, all group numbers and explanation of significant values are presented within the figure legends. Experiments were repeated twice to verify results.

### Data availability

All datasets generated during and/or analyzed during the current study are available from the corresponding author on reasonable request.

All schematics and models were created using Biorender.com.

## EXTENDED DATA FIGURES

**Extended Data Fig.1.**
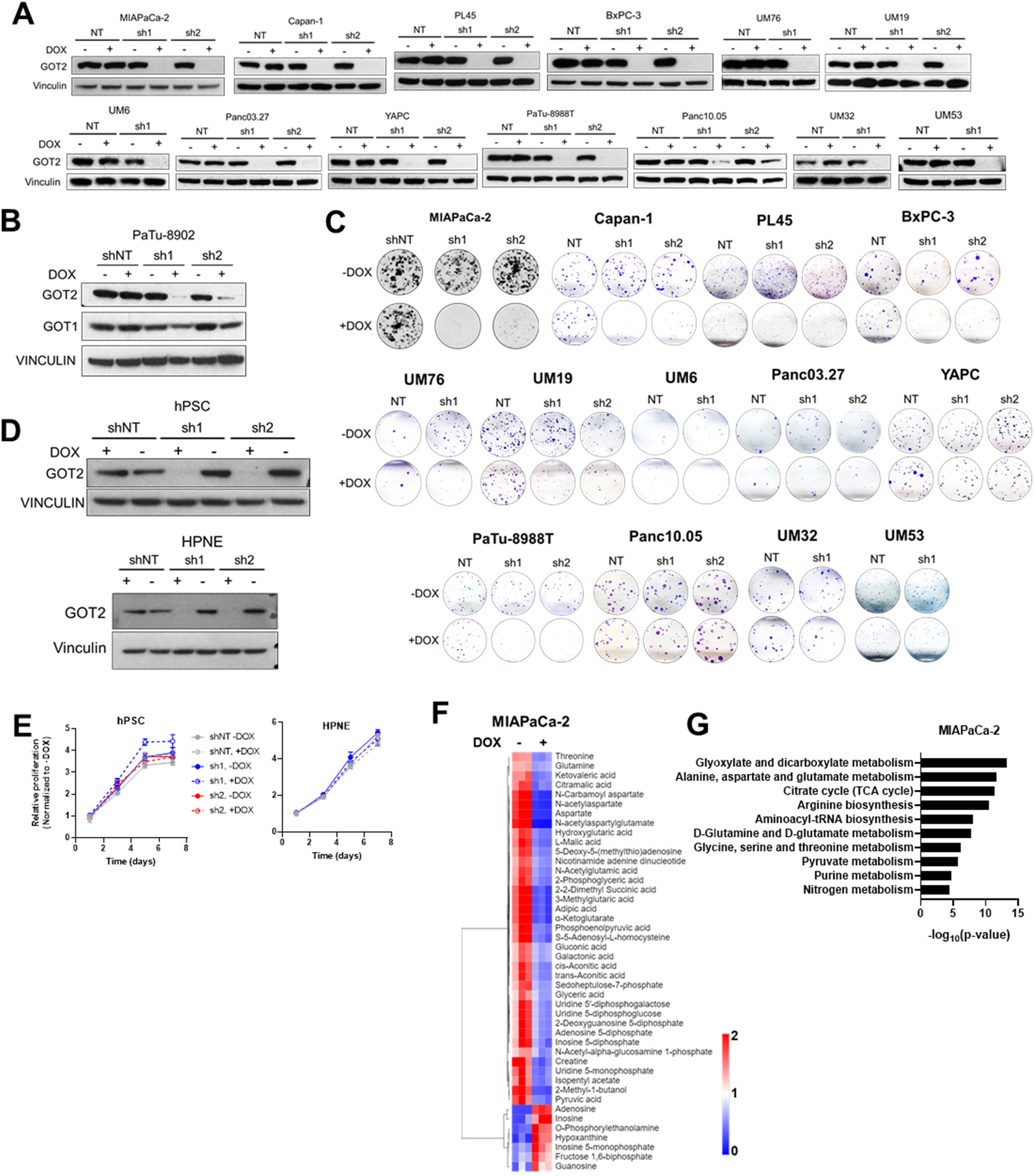
**A)** Western blots of GOT2 expression with Vinculin loading control in PDA cells expressing iDox-shNT, shGOT2.1, or shGOT2.2. **B)** Western blot for GOT1 and GOT2 with Vinculin loading control in PaTu-8902 iDox-shGOT2.1 cells. **C)** Images of representative wells showing colony formation in a panel of PDA cells expressing iDox-shNT, shGOT2.1, or shGOT2.2. **D)** Western blots of GOT2 expression with Vinculin loading control in HPNE or hPSC cell lines expressing iDox-shNT, shGOT2.1, or shGOT2.2. **E)** Relative proliferation of HPNE or hPSC cell lines expressing iDox-shNT, shGOT2.1, or shGOT2.2. **F,G)** Metabolites significantly changed between +DOX (n=3) and –DOX (n=3) (p<0.05, −1>log2FC>1) in MIAPaCa-2 iDox-shGOT2.1 cells as assessed by LC/MS. **F)** Heatmap depicting changes in relative metabolite abundances with 2D unsupervised hierarchical clustering. **G)** Metabolic pathways significantly changed, as determined via Metaboanalyst. Bars represent mean ± SD, *p<0.05, **p<0.01, ***p<0.001, ****p<0.0001.

**Extended Data Fig.2.**
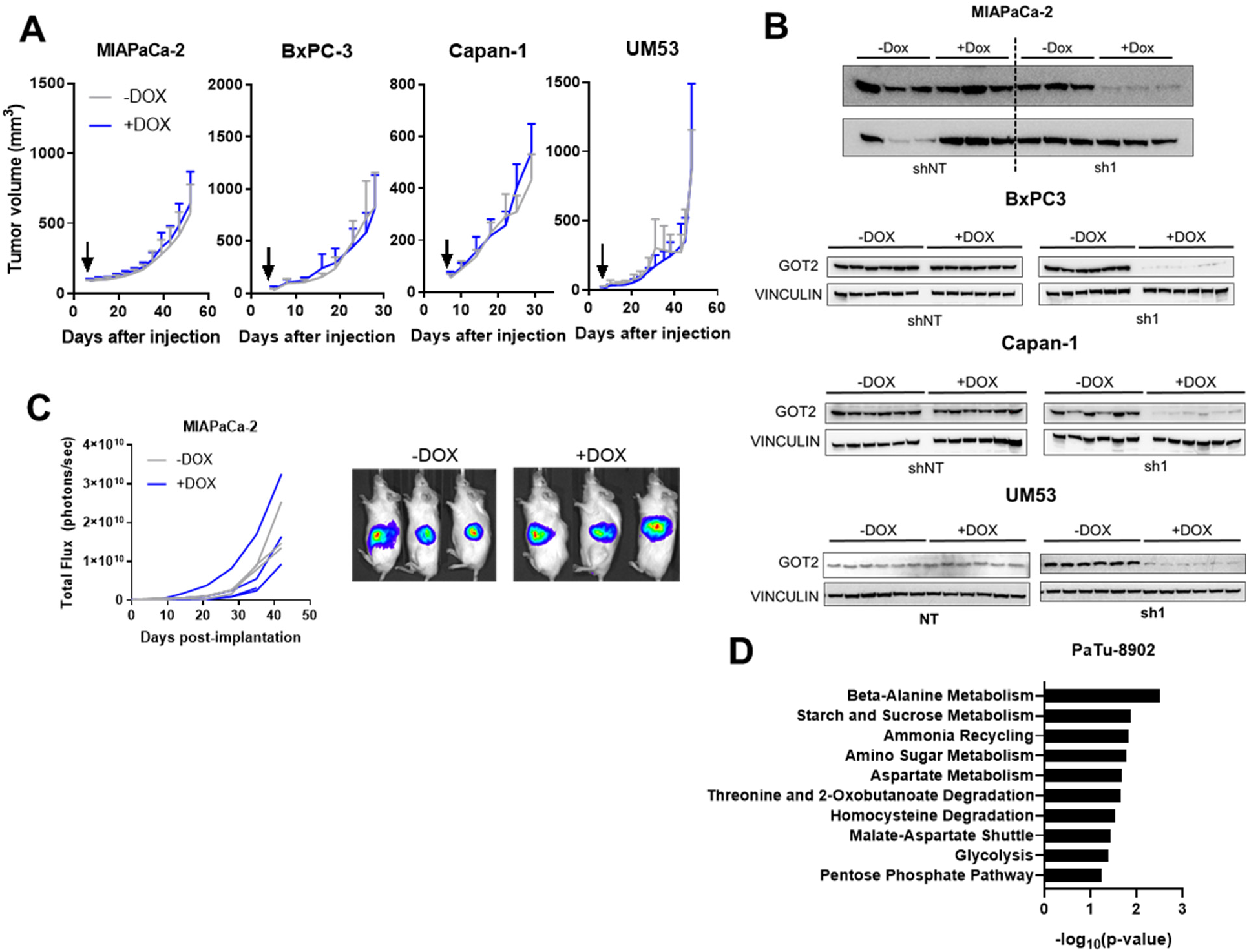
**A)** Growth across a panel of subcutaneous PDA control iDox-shNT subcutaneous xenografts in NSG mice (n=6 tumors per group). Arrows indicate administration of doxycycline chow. **B)** Western blot for expression of GOT2 and Vinculin loading control in PDA iDox-shNT or shGOT2.1 subcutaneous xenografts. **C)** Radiance of MIAPaCa-2 iDox-shGOT2.1 orthotopic tumors in NSG mice, with bioluminescent images at endpoint. **D)** Metabolic pathways significantly changed between +DOX (n=6) and –DOX (n=6) (p<0.05, −0.5>log2FC>0.5) in PaTu-8902 iDox-shGOT2.1 subcutaneous xenografts, as determined via Metaboanalyst. Bars represent mean ± SD.

**Extended Data Fig.3.**
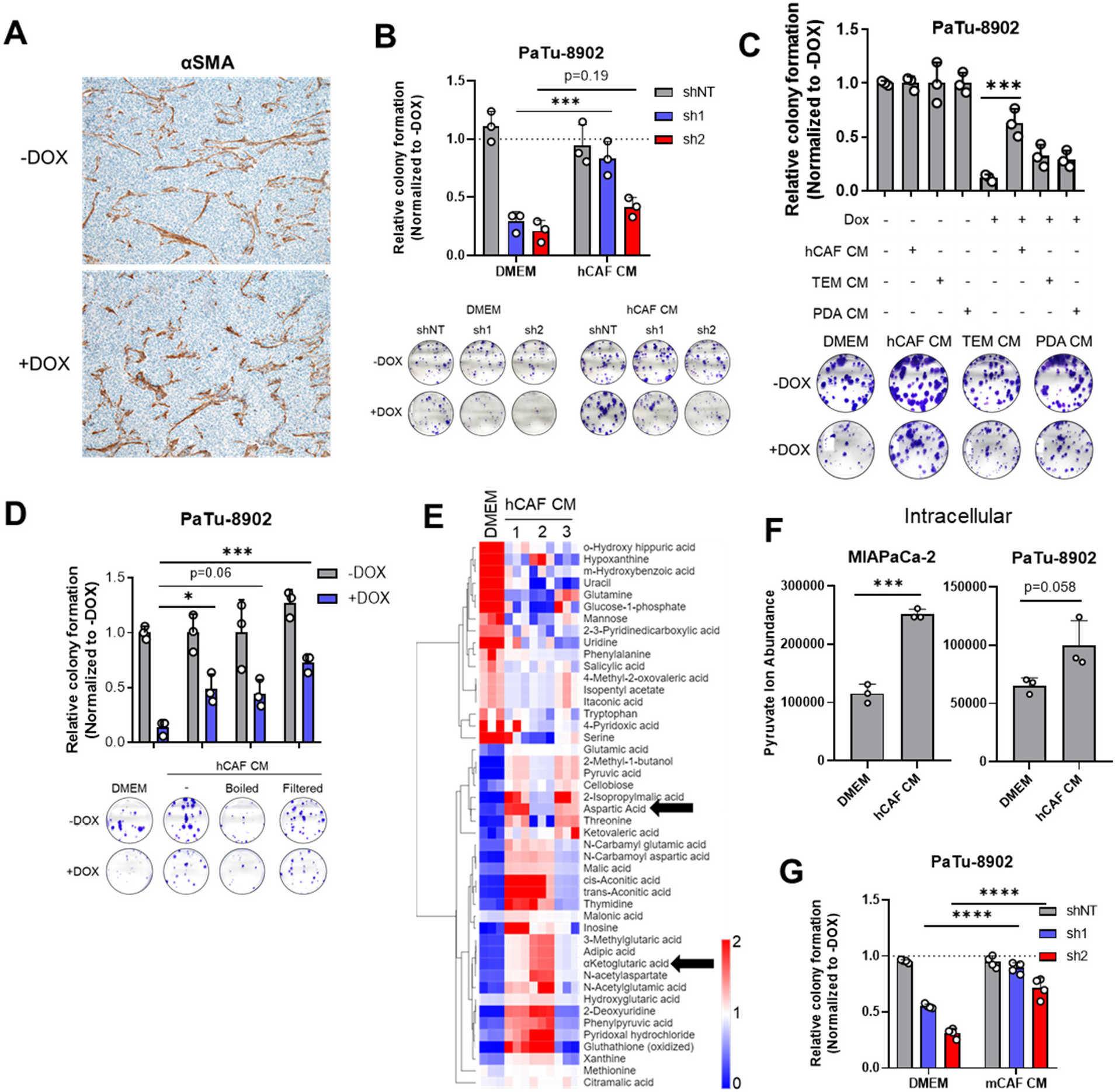
**A)** Representative αSMA staining in tissue slices from Fig.2A. **B)** Relative colony formation of PaTu-8902 iDox-shNT, shGOT2.1, or shGOT2.2 cells cultured in DMEM or hCAF CM, with images of representative wells. **C)** Relative colony formation of PaTu-8902 iDox-shGOT2.1 cells cultured in DMEM or hCAF CM, tumor educated macrophage (TEM) CM, or PaTu-8902 CM. **D)** Relative colony formation of PA-TU8902 iDox-shGOT2.1 cells cultured in DMEM or fresh hCAF CM, boiled CM, CM passed through a 3 kD filter, or CM subjected to freeze/thaw cycles, with images of representative wells. **E)** Relative abundances of metabolites significantly (p<0.05) changed between hCAF CM and DMEM, as determined by metabolomics. Asp and αKG are indicated with black arrows. **F)** Relative intracellular abundance of pyruvate in PDA cells cultured in DMEM or hCAF CM. **G)** Relative growth of PaTu-8902 iDox-shNT, shGOT2.1, or shGOT2.2 cells in DMEM, 1 mM Pyruvate, or mCAF CM. Bars represent mean ± SD, *p<0.05, **p<0.01, ***p<0.001, ****p<0.0001.

**Extended Data Fig.4.**
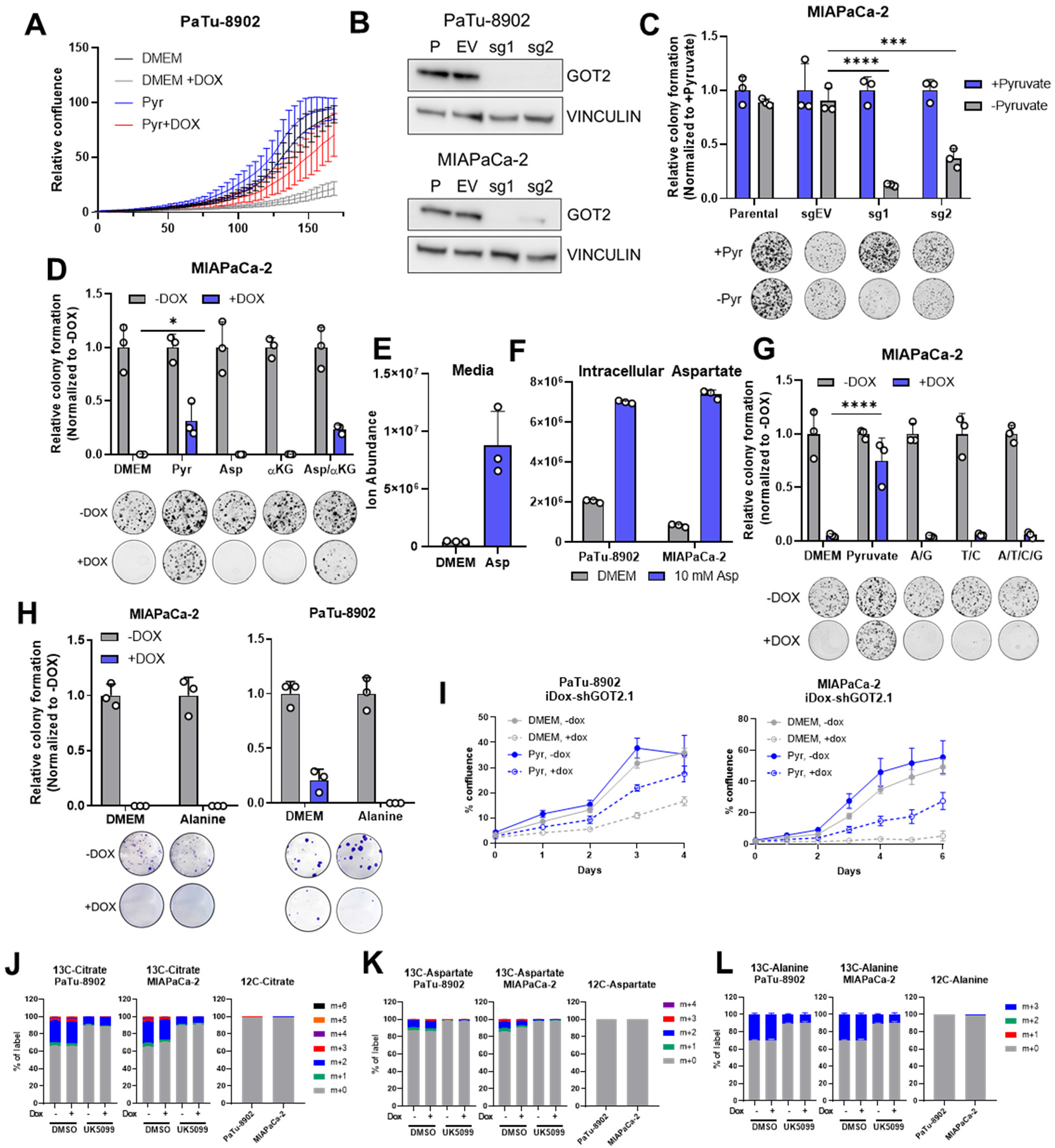
**A)** Relative confluence over time of PaTu-8902 iDox-shGOT2.1 cells cultured in DMEM or 1 mM pyruvate. **B)** Western blot for GOT2 expression in parental PDA cells, or cells expressing sgEV, sgGOT2.1, or sgGOT2.2, with Vinculin loading control. **C)** Relative colony formation of MIAPaCa-2 GOT2 KO cultured in DMEM or 1 mM pyruvate, with images of representative wells. **D)** Relative colony formation of MIAPaCa-2 iDox-shGOT2.1 cells cultured in DMEM, 10 mM aspartate, 4 mM αKG, or both Asp/αKG, with images of representative wells. **E)** Relative abundance of aspartate in DMEM or media containing 10 mM aspartate, as measured by LC/MS. **F)** Relative abundance of aspartate in PaTu-8902 or MIAPaCa-2 cells after culture in DMEM or 10 mM aspartate, as measured by LC/MS. **G)** Relative colony formation of MIAPaCa-2 iDox-shGOT2.1 cells cultured in DMEM or 100 μM of the indicated combinations of adenine (A), guanine (G), thymine (T), and cytosine (C), with images of representative wells. **H)** Relative colony formation of PaTu-8902 or MIAPaCa-2 iDox-shGOT2.1 cells cultured in DMEM or 1 mM alanine, with images of representative wells. **I)** Growth curves of PaTu-8902 or MIAPaCa-2 iDox-shGOT2.1 cells in media containing 1 mM unlabelled glucose and 1 mM U13C-Pyruvate. **J-K)** Labelling patterns of intracellular citrate (J), aspartate (K), or alanine (L) in PaTu-8902 or MIAPaCa-2 iDox-shGOT2.1 cells and treated with UK5099. Bars represent mean ± SD, *p<0.05, **p<0.01, ***p<0.001, ****p<0.0001.

**Extended Data Fig.5.**
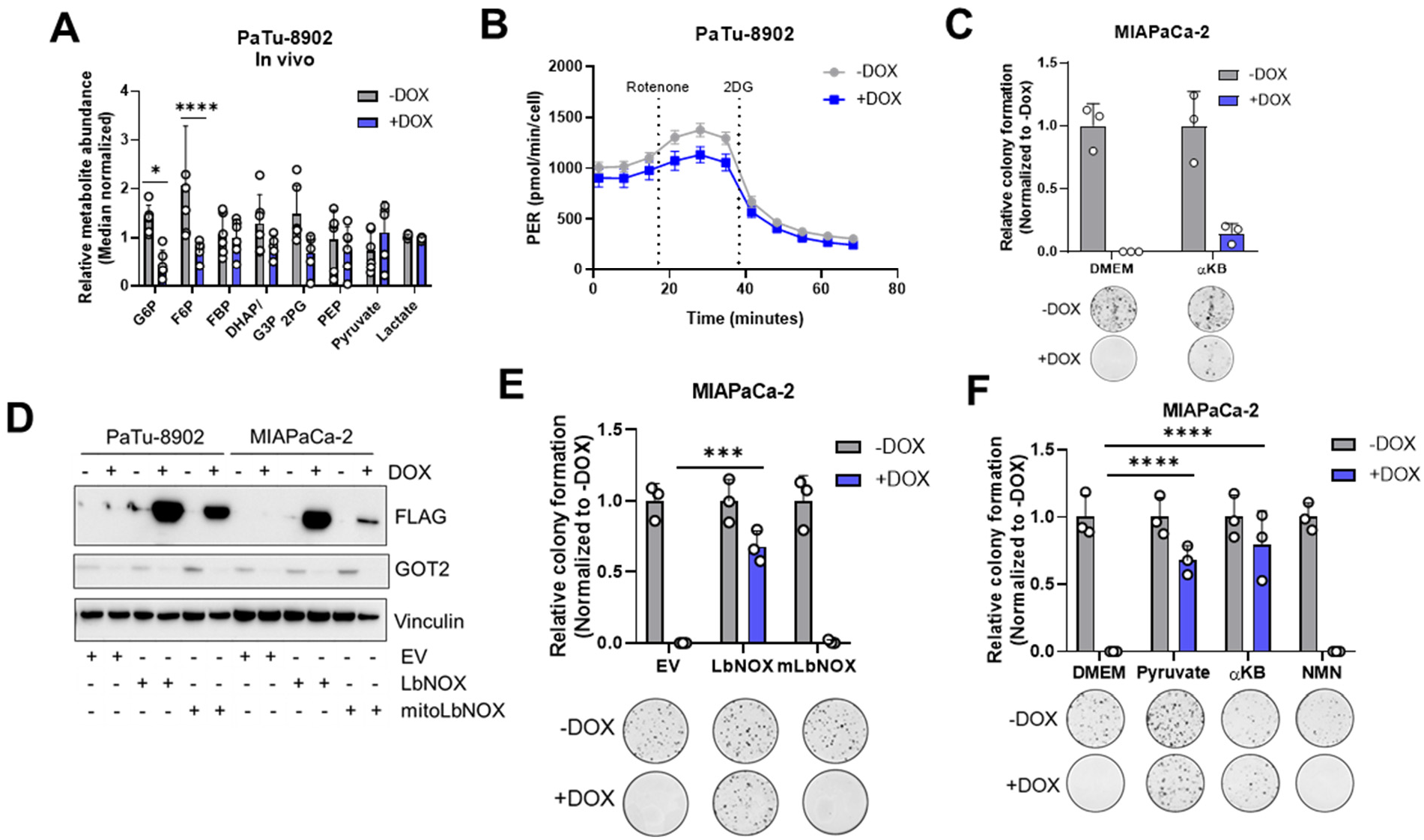
**A)** Relative abundances of glycolytic intermediates in PaTu-8902 iDox-shGOT2.1 subcutaneous xenografts. **B)** Proton efflux rate (PER) of PaTu-8902 iDox-shGOT2.1 cells cultured +DOX for 3 days, as determined by the Glycolytic Rate Assay. **C)** Relative colony formation of MIAPaCa-2 iDox-shGOT2.1 cells cultured in DMEM or 1 mM αKB, with images of representative wells. **D)** Western blot for Flag-tagged, doxycycline-inducible LbNOX or GOT2, with Vinculin loading control, in PaTu-8902 or MIAPaCa-2 iDox-shGOT2.1 cells. **E)** Relative colony formation of MIAPaCa-2 iDox-shGOT2.1 cells expressing cytosolic or mitochondrial LbNOX. **F)** Relative colony formation of MIAPaCa-2 iDox-shGOT2.1 cells cultured in DMEM, 1 mM pyruvate, αKB, or NMN, with images of representative wells. Bars represent mean ± SD, *p<0.05, **p<0.01, ***p<0.001, ****p<0.0001.

**Extended Data Fig.6.**
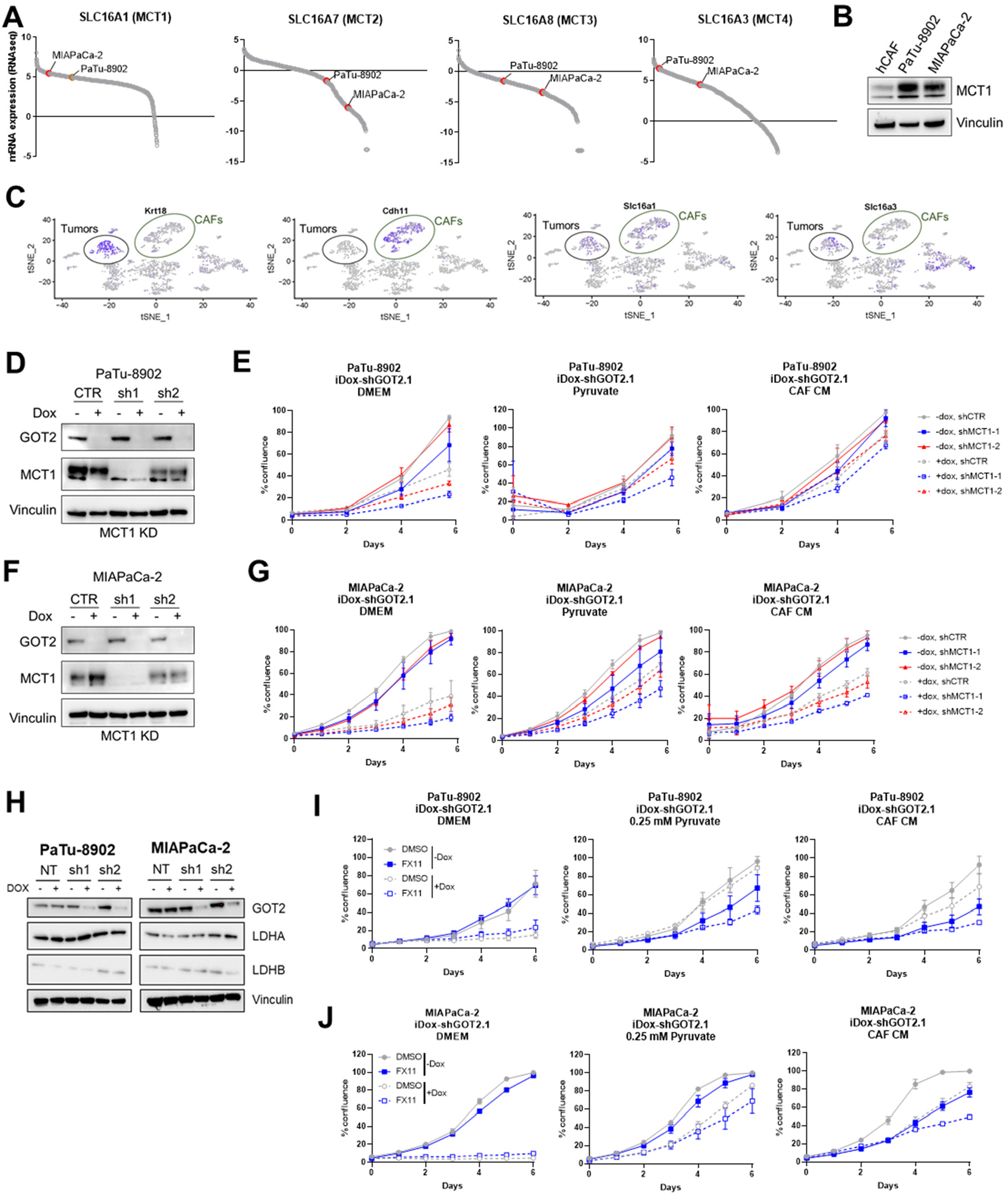
**A)** CCLE relative transcript expression levels of MCT family members in PDA cell lines. **B)** Western blot for MCT1, with vinculin loading control, in hCAFs or PaTu-8902 and MIAPaCa-2 cells. **C)** Single-cell RNA sequencing data from KPC syngeneic orthotopic tumors showing expression of Slc16a1 (MCT1) and Slc16a3 (MCT4) in CAF (CDH11) and epithelial (Krt18) populations. **D, F)** Western blots validating iDox-shGOT2.1 and constitutive MCT1 KD PaTu-8902 or MIAPaCa-2 cell lines, with vinculin loading controls. **E,G**) Growth curves of PaTu-8902 or MIAPaCa-2 iDox-shGOT2.1, MCT1 KD cell lines cultured in DMEM, 0.25 mM pyruvate, or hCAF CM. **H)** Western blots for LDHA and LDHB expression PaTu-8902 or MIAPaCa-2 iDox-shGOT2.1 cell lines, with vinculin loading controls. **I,J)** Growth curves of PaTu-8902 or MIAPaCa-2 iDox-shGOT2.1 cell lines cultured in DMEM, 0.25 mM pyruvate, or hCAF CM and treated with 25 μM FX11.

**Extended Data Fig.7.**
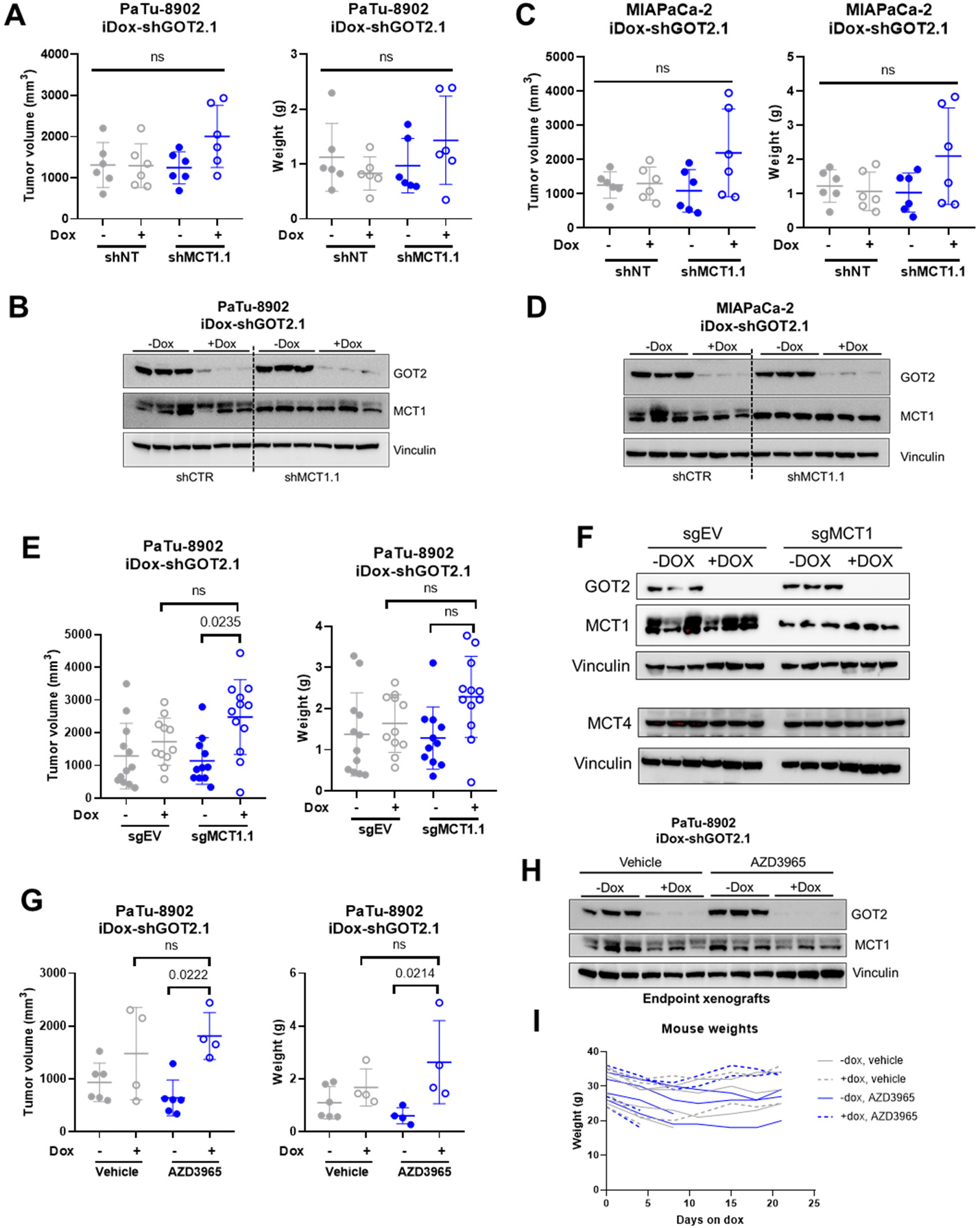
**A,C)** Graphs depicting endpoint tumor volumes (left) or tumor weights (right) of PaTu-8902 or MIAPaCa-2 iDox-shGOT2.1, MCT1.1 KD subcutaneous xenografts in NSG mice. **B,D)** Western blots confirming GOT2 KD and MCT1 KD, with vinculin loading controls, with protein lysates from xenografts in A) and C). **E)** Graphs depicting endpoint tumor volumes (left) or tumor weights (right) of PaTu-8902 iDox-shGOT2.1, sgMCT1.1 subcutaneous xenografts in NSG mice. **F)** Western blots confirming GOT2 KD and MCT1 KO along with MCT4 expression, with vinculin loading controls, with protein lysates from xenografts in E). **G)** Graphs depicting endpoint tumor volumes (left) or tumor weights (right) of PaTu-8902 iDox-shGOT2.1 subcutaneous xenografts treated with 100 mg/kg AZD3965 via daily oral gavage. **H)** Western blots confirming GOT2 KD and showing MCT1 expression, with vinculin loading controls, with protein lysates from xenografts in G). **I)** Weights of mice fed normal or doxycycline chow and treated with vehicle or AZD3965.

**Extended Data Fig.8.**
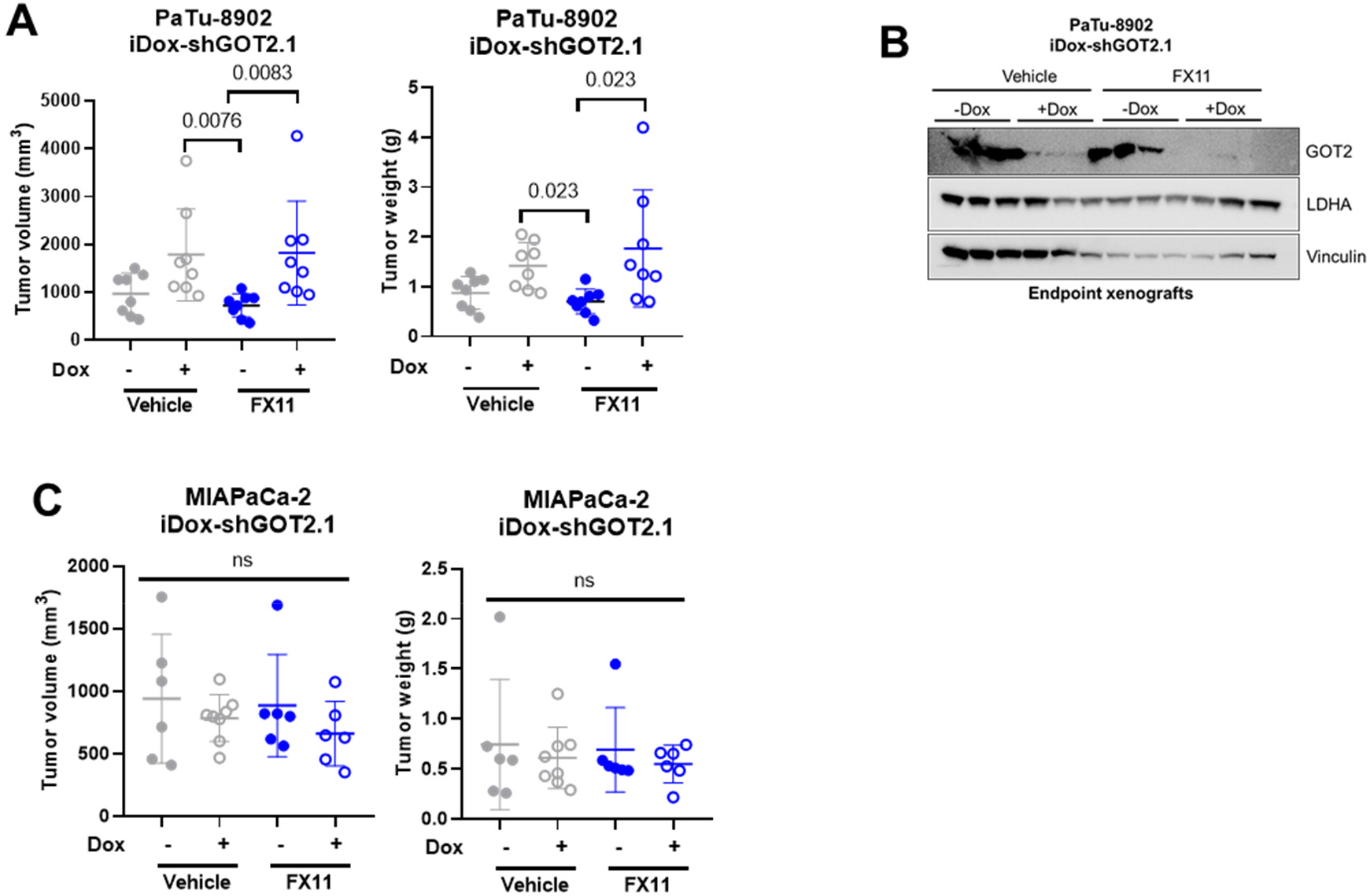
**A,C)** Graphs depicting endpoint tumor volumes (left) or tumor weights (right) of PaTu-8902 or MIAPaCa-2 iDox-shGOT2.1 subcutaneous xenografts treated with 2 mg/kg FX11. **B,D)** Western blots confirming GOT2 KD and showing LDHA expression, with vinculin loading controls, with protein lysates from xenografts in A) and C).

**Extended Data Fig.9.**
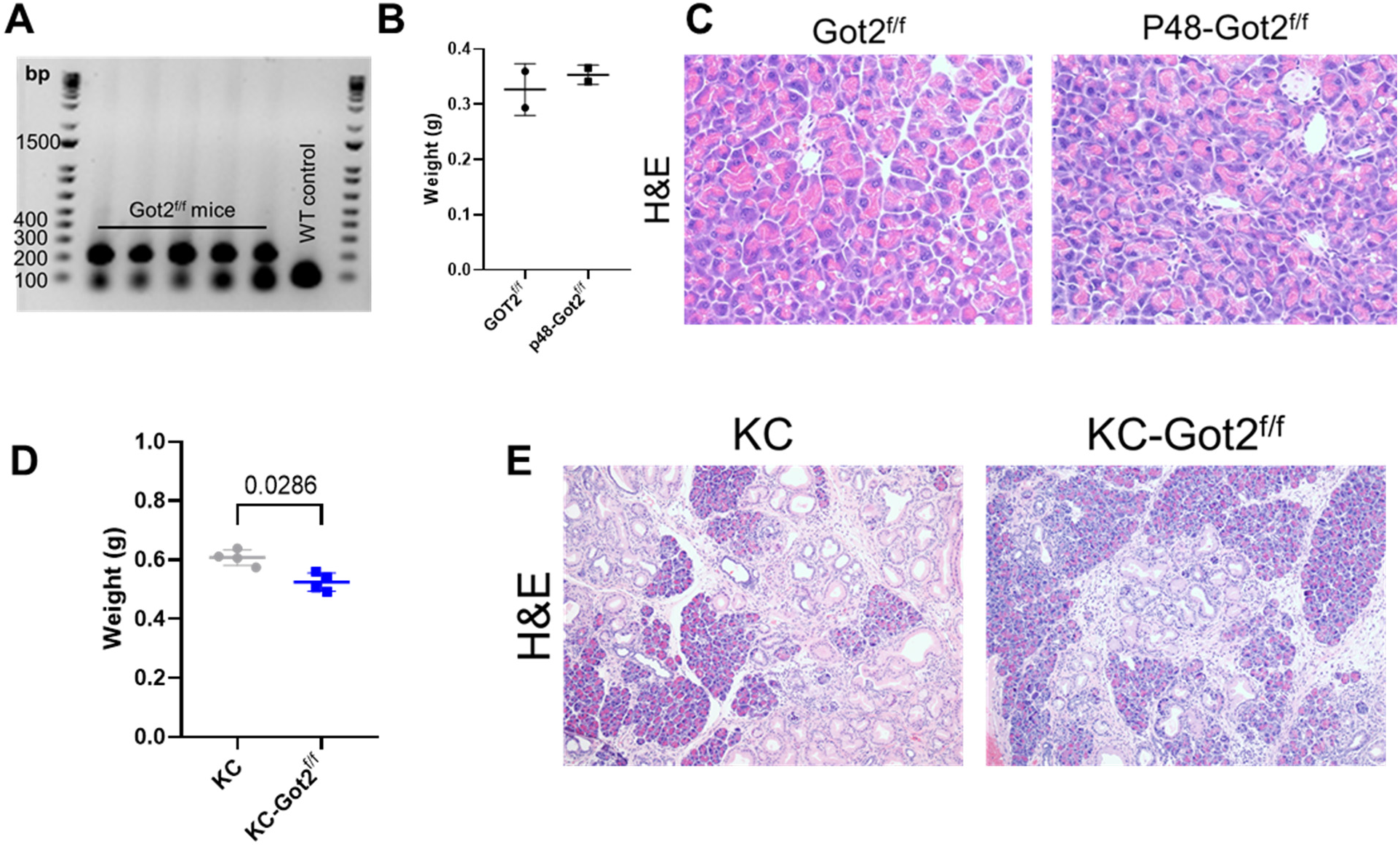
**A)** DNA gel showing representative genotyping of DNA isolated from tails of Got2^f/f^ mice or wild type (WT) control. **B)** Weights of healthy pancreata from 3 month old wild type or Ptf1a-Got2 mice. **C)** Representative H&E images of pancreata from 3 month old wild type or Ptf1a-Got2 mice, 10x. **D)** Weights of pancreata from 6 month old KC or KC-Got2 mice. **E)** Representative H&E images of pancreata from 6 month old KC or KC-Got2 mice, 10x.

